# Conserved emergent traits enable phylogenetic prediction of community function

**DOI:** 10.64898/2026.06.09.731074

**Authors:** Uroš Gojković, Zorana Miloradović, Nikola Popović, Goran Vukotić, Nina Medaković, Nemanja Stanisavljević, Nemanja Kljajević, Djordje Bajić

## Abstract

Predicting the function of a microbial community from the identity of its members is a key goal of ecology and biotechnology. Here, we show that the systematic variation in how strains contribute to community function is phylogenetically conserved. This allows us to predict the collective function of communities solely from their member’s taxonomic identity. Using soymilk fermentation as a model, we measured three industrially relevant functions − acidification, texture, sensory grade − across 307 synthetic communities combining 33 phylogenetically diverse lactic acid bacteria strains. The functional effect of a strain scaled linearly with the function of the receiving community, allowing us to summarize each strain’s contribution across communities with two parameters, intercept and slope. These parameters were phylogenetically conserved, allowing us to impute their values for 17 previously unassayed strains, and use them to predict the three functions in 14 newly assembled communities. By redefining community function in terms of conserved emergent species traits, our results enable predictive engineering of microbial consortia at scale, and open a path towards genomic and evolutionary dissection of complex community phenotypes.

## Introduction

Modern microbial biotechnology relies on culture collections of remarkable scale. Public microbial repositories archive a substantial fraction of cultivable microbial diversity (*1,2*), and industrial biobanks routinely hold thousands of strains, characterized to a variable degree (*3–5*). These resources are increasingly being mobilized not only to select individual production strains, but to assemble synthetic consortia (*6,7*) whose collective metabolism can unlock routes inaccessible to monocultures (*8–12*), broadening the range of usable substrates (*13*) and produced compounds(*14*). With potential applications spanning pharmaceutical production (*15,16*), biofuel synthesis (*17,18*), waste valorization (*19,20*) and sustainable agriculture (*21,22*), we urgently need strategies to rationally leverage microbial biobanks for the engineering of functional communities with minimal experimental and economic investment (*23,24*).

From a fundamental standpoint, connecting the identity of individual species to community-level outcomes is a major goal in ecology (*25–27*). Unlike single strains, whose phenotypes can increasingly be inferred from their genome (*28–30*), the function of multi-species communities emerges from a complex network of interactions between its members (*31–33*), which makes it much more challenging to predict. Several complementary approaches have been developed to navigate this complexity. Mechanistic genome-scale models, together with their community-level extensions (*34–37*), can be used to predict function from first metabolic principles, but high quality predictions still depend on strain-specific curation, and experimental constraints that are difficult to obtain at scale. High-throughput trial-and-error screens can bypass this issue, but are limited by combinatorial explosion, as even modest sized collections rapidly exceed the number of consortia that can be experimentally assembled (*38*).

Statistical approaches such as fractional-factorial design of experiments combined with regression (*39–42*) can predict the function of untested combinations from a training set, but they assume that interactions remain sparse and landscapes are smooth (*43*), and their predictions are confined to the strains experimentally probed during training. More recently, a number of studies have shown that it is possible to connect the genetic content of a community to functional outcomes using regression or machine-learning approaches (*44–47*). However, these strategies require fully sequenced and annotated genomes and either a substantial mechanistic understanding of the function of interest (*45*), or large amounts of training data (*44*). Given the growing scale of modern biobanks, to what extent is it possible to predict the function of a community with minimal information about its members − ideally, just their taxonomic identity?

To answer this question, we hypothesized that a conserved high-level descriptor might exist that is able to predict a strain’s functional effect when combined with other strains, i.e. across “community backgrounds”. If such a descriptor were conserved, it could be inferred from a strain’s taxonomy (e.g. 16S ribosomal RNA sequence), allowing us to infer the strain’s functional contribution even in absence of any direct functional measurement. A candidate for this descriptor has recently emerged from work on community function landscapes, maps between community structure and a function of interest. In particular, the functional effect of adding a focal species *i* to a background community has **s** been shown to often depend linearly on the function of that background, a relationship referred to as a “functional effect equation” (FEE) (*31,48*), with the form *ΔF*_*i*_*(s)* = *a*_*i*_ + *b*_*i*_*F(s)* + *θ*_*i*_*(s)*,, where Δ***F***_***i***_(s) = ***F***(s + *i)* – ***F***(s) is the change in community function caused by the addition of species *i*, is the function of the background community before the addition (Fig. 1, Methods). This linear functional effect equation summarizes the functional contribution of a species with only two parameters, the intercept *a*_*i*_ and slope *b*_*i*_, and it has been used to predict community function with notable accuracy (*48,49*). Functional effect equations, conceptually analogous to global epistasis patterns observed in quantitative genetics (*50–52*), have been observed in community-function landscapes across phylogenetic groups and functions, ranging from bacterial starch degradation to yeast sugar consumption and plant community biomass (*48,49*). If these FEE slopes and intercepts were themselves predictable from a species taxonomic identity, then we could impute their values for untested strains, extending the predictive capability of the method potentially by orders of magnitude, from the size of the tested combinatorial space to the phylogenetic coverage of the entire collection at hand.

**Figure 1.**
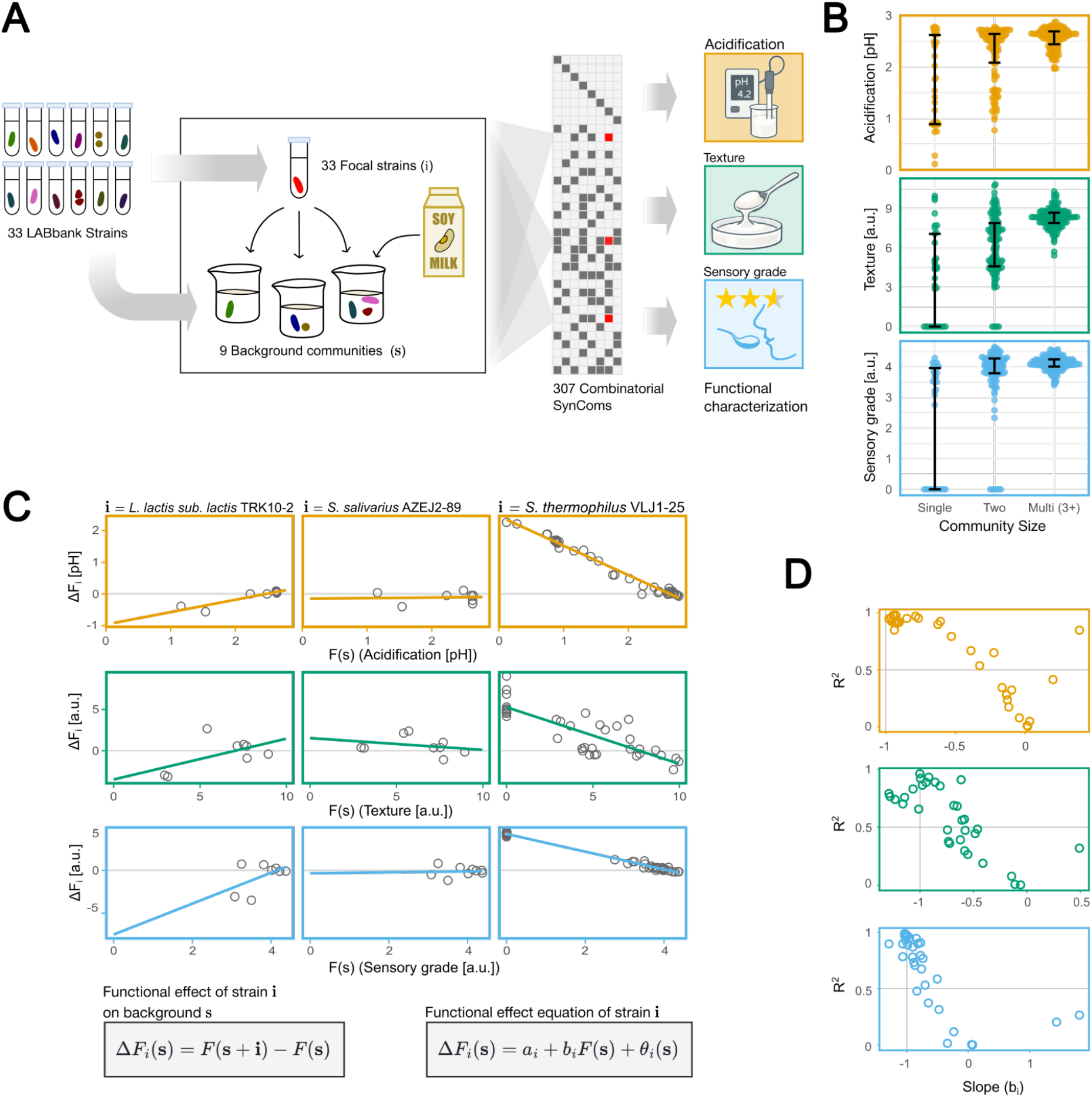
**A)** Schematic overview of the experimental setup. Using 33 strains from the LABbank collection, we have constructed 307 combinatorial synthetic communities (SynComs). Each of these SynComs was then inoculated in soymilk and, upon fermentation, three functional parameters were measured: acidification (yellow), texture (green) and sensory grade (blue). **B)** Distribution of functional outputs across community sizes. Each dot represents one of 307 tested combinations. Black bars indicate median ± interquartile range. Consortia composed of two or more members outperformed the monocultures across all three functions (t-test p<0.05, for all three functions). **C)** Functional effect equations for three representative species across the three measured functional outputs (Supplementary Fig. S1 - S3 for complete data visualization). Each point represents the difference in function when the given species is added to the background. Ordinary least-squares linear fits were used to estimate functional effect equations, characterized by a slope *b* and intercept *a*. **D)** Relationship between slope (b) and R^2^ for all 33 species across the three measured functions. Each dot represents one species. A large majority of strains in all three functions deviate from the expectations under either an additive *(b*=0) or a random landscape*(b*=-1, and R ^2^ = 0.5).

Here, we test this hypothesis in a large lactic acid bacteria (LAB) biobank (*53*) in the context of soymilk fermentation. Our experiments encompass three industrially relevant functional readouts of qualitatively different levels: a metabolic phenotype (acidification), a physical phenotype (texture), and an integrated organoleptic phenotype (sensory grade). We show that global epistasis governs all three functions, with FEEs providing moderate to excellent predictive accuracy. We then show that the FEE parameters *a*_*i*_ and *b*_*i*_ are largely conserved, and use this conservation to predict the function of consortia composed of strains entirely absent from the training set, using only their 16S rRNA gene sequence derived identity. These results establish FEEs as a practical tool for biobank-scale design of microbial consortia. From a fundamental perspective, the demonstration that FEE parameters are heritable suggests they can be treated as bona fide ecological traits, opening the prospect of mapping the genomic determinants of collective community function.

## Results

### Global epistasis governs three classes of soymilk fermentation function

We first sought to establish whether FEEs describe how individual strains contribute to relevant community functions in soymilk fermentation. To that end, we drew on the LABbank collection, a biobank of over 5000 lactic acid bacteria assembled over several decades from traditional fermented dairy products, fermented plant-based products (e.g., sauerkraut, alfalfa silage), and rumen content (*53*). From this repository we selected 33 strains spanning nine genera (Methods), chosen to maximize phylogenetic breadth including both dairy- and plant-origin lineages (Fig. 1A). From these strains, we assembled 307 combinatorial synthetic communities (SynComs) of varying richness (Methods), inoculated each in soymilk, and quantified the three functional readouts: acidification (measured as difference in pH after 15 hours of fermentation), texture (measured as gel hardness), and sensory grade (panel-scored) (Methods, Fig. 1A, Supplementary table 1).

A first look at the data confirms that consortium-level optima exist beyond the reach of single strains for all three functions (t-test comparing single versus multi-species consortia, p<0.05 for all three functions; Fig. 1B), motivating a quantitative description of how strains functionally combine. For each function, we measured the functional effect observed upon the addition of a strain *i* to different background communities ***s s*, Δ*F***_***i***_**(s)**, and regressed this against the function of the background *F(***s**) . For the majority of strains across all three functions, **Δ*F***_***i***_**(s)** depended approximately linearly on *F*(s) (Fig. 1C, Fig. S1-3). Across the panel, FEEs explained a large fraction of the variance in all three functions (median R^2^ = 0.90, 0.66, 0.78 for acidification, texture, and sensory grade, respectively), suggesting they largely summarize the functional effects of most of our strains.

The three values describing the FEE 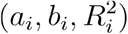 can be useful to gauge the characteristics of the underlying functional landscapes (*54*). In particular, a purely additive landscape, in which a strain’s contribution is independent of community background, is expected to trivially produce FEEs with slopes ***b***_***i***_ ≈ 0. A purely random landscape, in which a strain’s effect is fully uncorrelated, produces slopes ***b***_***i***_ ≈ − and 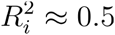 as a consequence of regression to the mean (*54*). The observed 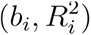 pairs across our 33 strains are generally inconsistent with either limit, and most species clearly deviate from both the random landscape expectation and from additivity in all three functions (Fig. 1D). This suggests that the FEEs capture the dominant patterns dictating how a strain contributes across communities, and might be predictive of its effects in experimentally unseen combinations.

### Global epistasis can predict the function of microbial communities

To test whether FEEs can be used for community-level prediction in our three functions, we applied the previously described “stitching” approach. In this method, the function of any consortium is predicted by iteratively adding species one at a time, using each species’ FEE to estimate its contribution to the current background function at each step (*48*). In iterated 10-fold cross-validation across the 307 Syncom panel, the FEE-based stitching method achieved high accuracy for acidification, and more moderate accuracies for texture and sensory grade (Fig. 2A, black circles; Methods).

**Figure 2.**
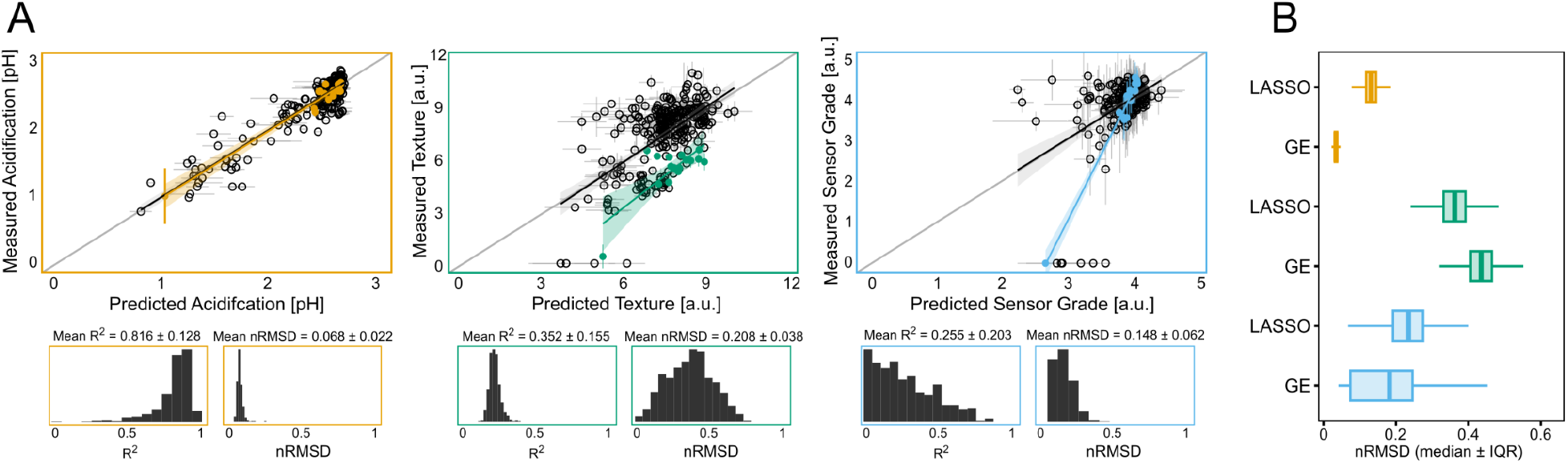
Functional effect equations predict three functional traits across SynComs. **A)** We used functional effect equations (FEEs) derived for 33 LAB strains to predict three community level functions utilizing the “stitching” approach (*48*). Empty circles represent predictions from iterated 10-fold cross-validation (N=500), while the coloured points correspond to 17 validation SynComs (Methods). Error bars indicate ±SD across cross-validation iterations and experimental measurements (note that acidification was measured once). The black lines with grey shading represent linear fits with 95% CI for the cross-validation iterations. Coloured lines (orange, green, and blue) correspond to linear fits with 95% CI for the 17 validation SynComs. Model performance was evaluated using R^2^ and normalized root mean square deviation (nRMSD), whose distribution for cross-validation iterations is shown in the bottom sub-panels. In the case of validation communities, predictions for acidification (orange) showed the best accuracy for both metrics (mean R^2^ = 0.816 and nRMSD = 0.068, N=17), while texture (green) and sensor grade (blue) showed lower levels of predictability. Texture exhibited a higher R ^2^ than sensor grade (0.352 vs 0.255, N=17) but also a higher prediction error (nRMSD = 0.208 vs 0.148, N=17). **B)** Comparison of predictive performance between the global epistasis model and LASSO regression assessed by bootstrapping (10000 iterations, Methods). Distribution of nRMSD are shown for acidification (orange), texture (green) and sensor grade (blue). Global epistasis consistently outperforms LASSO for acidification and sensor grade, while LASSO shows a slightly lower prediction error for texture (paired t-test, p<10^−16^, p=1, p<10^−16^, for acidification, texture and sensory grade, respectively). See Supplementary Fig. S4 for a more detailed comparison.

Is it possible to generalize these predictions to communities not seen during FEE estimation? Using the FEEs fit on the full 307 Syncom panel, we generated predictions for all untested 2-, 3-, and 4-member communities, then experimentally validated 17 representative combinations spanning the full predicted range of function (Methods). Prediction performance for these communities broadly matched the cross-validation results (Fig. 2A, coloured points). Rank predictions were very strong and significant for acidification and sensory grade, but non-significantly correlated for texture (Spearman’s ρ = 0.77, 0.32, and 0.86; p = 3.39 × 10^−4^, 0.21, and 1.21 × 10^−5^, for acidification, texture, and sensory grade, respectively; Fig. 2A). The model is thus able to quantitatively distinguish high from low-performing communities in 2 out of 3 functional readouts, even in presence of experimental biases (e.g. batch effects), which is relevant in applied contexts.

To benchmark this approach against a standard alternative, we trained a LASSO regression model with explicit pairwise interaction terms on the same 307 Syncom training set, following the approach previously applied to community function prediction (*55*). The FEE-stitching method outperformed LASSO for acidification (nRMSD 0.04 vs 0.13) and sensory grade (0.19 vs 0.26). For texture, the result was mixed: FEEs captured a substantially larger fraction of variance (R^2^ 0.57 vs 0.27) but with a somewhat higher prediction error (nRMSD 0.44 vs 0.36; see also Supplement Fig. S4A-B). Because the comparison rested on only 17 validation communities, we used bootstrapping to verify that these performance differences were robust (Methods). The bootstrap distributions confirmed the FEE outperformed LASSO for acidification and sensory grade, and at least matched it for texture (paired t-test, p<10^-16^, p=1, p<10^-16^, for acidification, texture and sensory grade, respectively, Fig. 2B). FEE-based prediction therefore matches or exceeds the performance of an established interaction-explicit regression method, with the advantage that they summarize each strain’s contribution into two interpretable parameters instead of estimating a separate coefficient for every possible interaction.

### Phylogenetic conservation of FEE parameters enables extrapolation to new strains

A closer inspection of the data suggested that FEE parameters in our dataset might be phylogenetically conserved (Fig. 3A). Thus, we hypothesized that FEE parameters *a*_*i*_ and *b*_*i*_ are themselves conserved and thus predictable from a strain’s phylogenetic identity. To test this hypothesis, we computed two complementary measures of phylogenetic signal, Pagel’s λ and Abouheif’s *C*_**mean**_, on the FEE parameters fit to the 33 strains (*56, 57*) (Methods).

**Figure 3.**
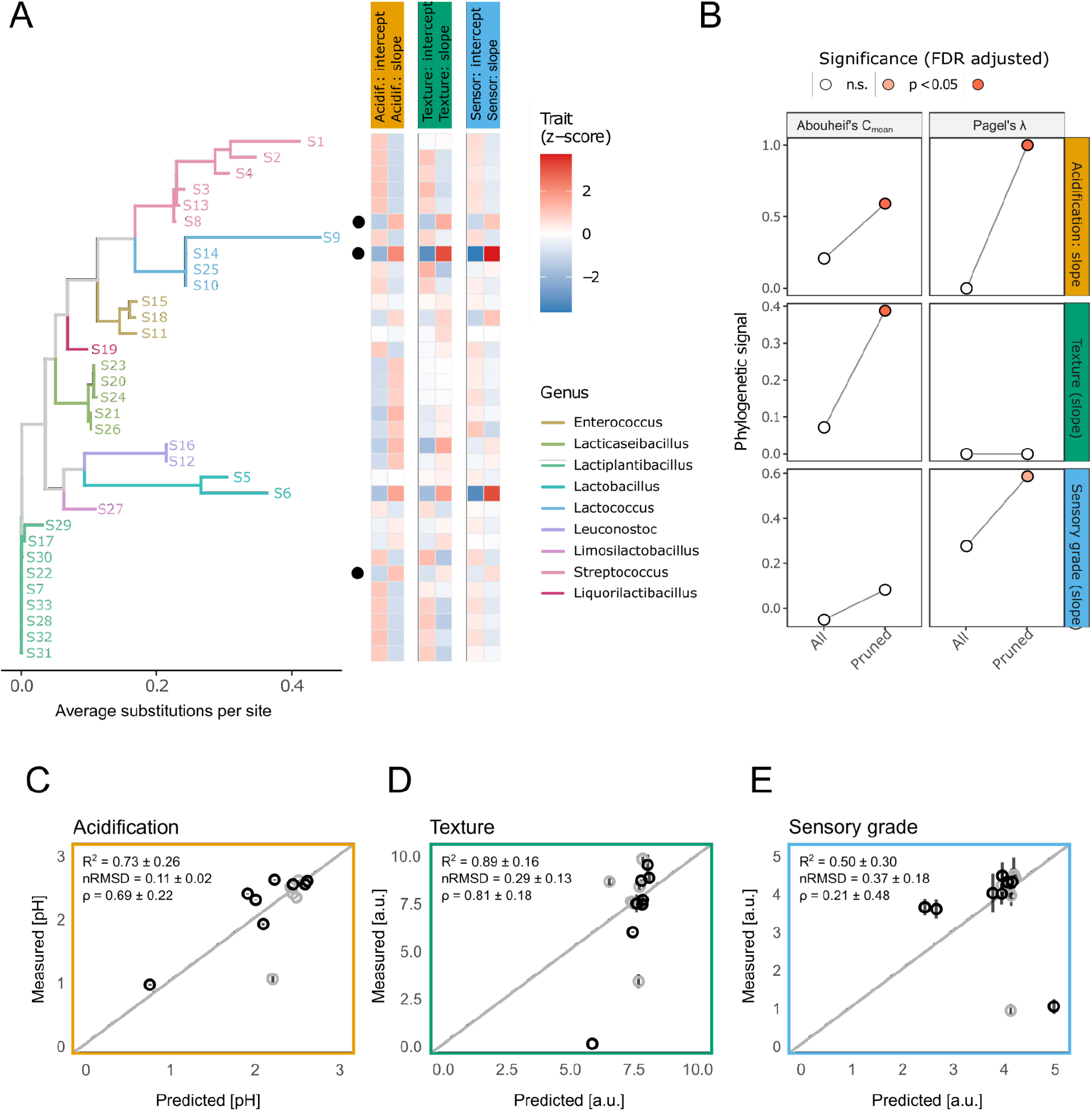
Phylogenetic conservation of FEE parameters predicts the function of SynComs composed of previously not assayed strains. **A)** Phylogenetic signal in FEE parameters (slopes and intercepts) across strains, in all three functions. **B)** Phylogenetic conservatism analysis of FEE slopes and intercepts. When all strains are included, signal is generally weak; however, leave-one-out analyses (Supplementary Figure S5 − S7) identified a small number of phylogenetically discordant isolates (specifically strains S14, S22, and S8) that strongly influence signal estimates. After excluding these strains (“Pruned” tree), descriptors show strong and significant phylogenetic signals across the majority of the tree. **C-E)**Prediction of community function for communities composed fully (black) or partially (grey) of previously unassayed strains. Predicted versus measured values for acidification (D) texture (E), and sensory quality (F). Predictions based solely on phylogenetic imputation capture a substantial fraction of the observed variation, with strongest performance for acidification and preservation of rank ordering across traits. Predictability metrics shown on the panels (R2, nRMSD, ρ) represent SynComs fully composed of unassayed strains (N=8, Supplementary table 2 for more exhaustive breakdown).

When applied to the full panel, the phylogenetic signals appeared non-significant for most descriptors. However, a leave-one-out influence analysis revealed three strains (S8, S14, and S22) whose FEE parameters diverged sharply from those of their closest phylogenetic relatives across all three phenotypes (Fig. 3B; Supplementary Figs. S5, S6), and whose inclusion was responsible for the lack of overall signal. Such discordance is expected if a number of strains sharply diverge from their phylogenetic neighbors, e.g. through horizontal gene transfer (*58*), but also from possible technical artifacts derived from e.g. contamination, that inevitably percolate in high-throughput experiments, and can propagate through the FEE estimation procedure. Given that the same strains disrupt conservation in all three functions, we believe the latter option to be the most likely in our data. After “pruning” the phylogenetic tree by excluding the three discordant isolates, a markedly different pattern emerged, with FEE parameters exhibiting strong and significant phylogenetic signals in all three functions (Fig. 3B).

While these observations indicate that FEE parameters behave as phylogenetically conserved traits across most of the biobank, in-sample identification of outliers cannot, by itself, validate this conclusion. To provide an unbiased test, we used the pruned phylogeny to impute FEE parameters for 17 additional strains drawn from the LABbank collection that had not been included in any of the 307 training SynComs (Methods, Supplementary Fig. S). We then experimentally assayed 14 communities either partially (N = 6) or fully (N = 8) composed of these previously unassayed strains, and compared the measured functions to predictions generated using phylogenetically imputed FEE parameters. The resulting predictions captured a substantial fraction of the observed functional variation (Fig. 3C–E).

Predictability for communities composed fully of previously unassayed strains was highest for acidification and texture, in both cases recovering the rank ordering of communities (Spearman’s ρ = 0.76 and 0.85, p=0.028 and 0.006, respectively; N=8). In the case of sensory grade, predictions were less well calibrated (Spearman’s ρ = 0.20, p=0.63, N=8, Supplementary Table 2), although the pattern seems driven by only a few outliers. Overall, this is to the best of our knowledge the first demonstration that a strain’s functional contribution across community contexts can be predicted from its 16S rRNA gene sequence-derived identity alone, with no prior mechanistic or phenotypic information.

### Predictability is controlled by functional interaction structure

The differential predictive accuracy observed in our results suggests that the underlying functional landscapes might have a fundamentally different structure. Previous theory suggested that complex functions, such as those requiring coordinated contributions from multiple species, give rise to rugged functional landscapes that are inherently less predictable (*55,59*). We therefore hypothesized that differences in predictability between our three functions reflect differences in the ruggedness of the underlying functional landscapes.

To explore this possibility, we reasoned that by definition, more rugged landscapes differ in the magnitude and order of interaction terms underlying each landscape. The absolute magnitude of interactions can therefore be used as a proxy measure of landscape ruggedness. To quantify the magnitude of these interactions, we focused on eight three-species subsets within our dataset for which the full combinatorial landscape was experimentally characterized, including all single strains, pairwise combinations, and the corresponding three-member communities (Fig 4A and Supplementary Fig. S9). Within these subsets, we quantified interactions as deviations of the co-culture function from the expectation under a Bliss null model of independence, which is appropriate for bounded functions (Methods). This analysis revealed that the texture function is shaped by substantially stronger interactions, both pairwise and third-order, compared to acidification and sensory quality (Fig. 4B).

**Figure 4.**
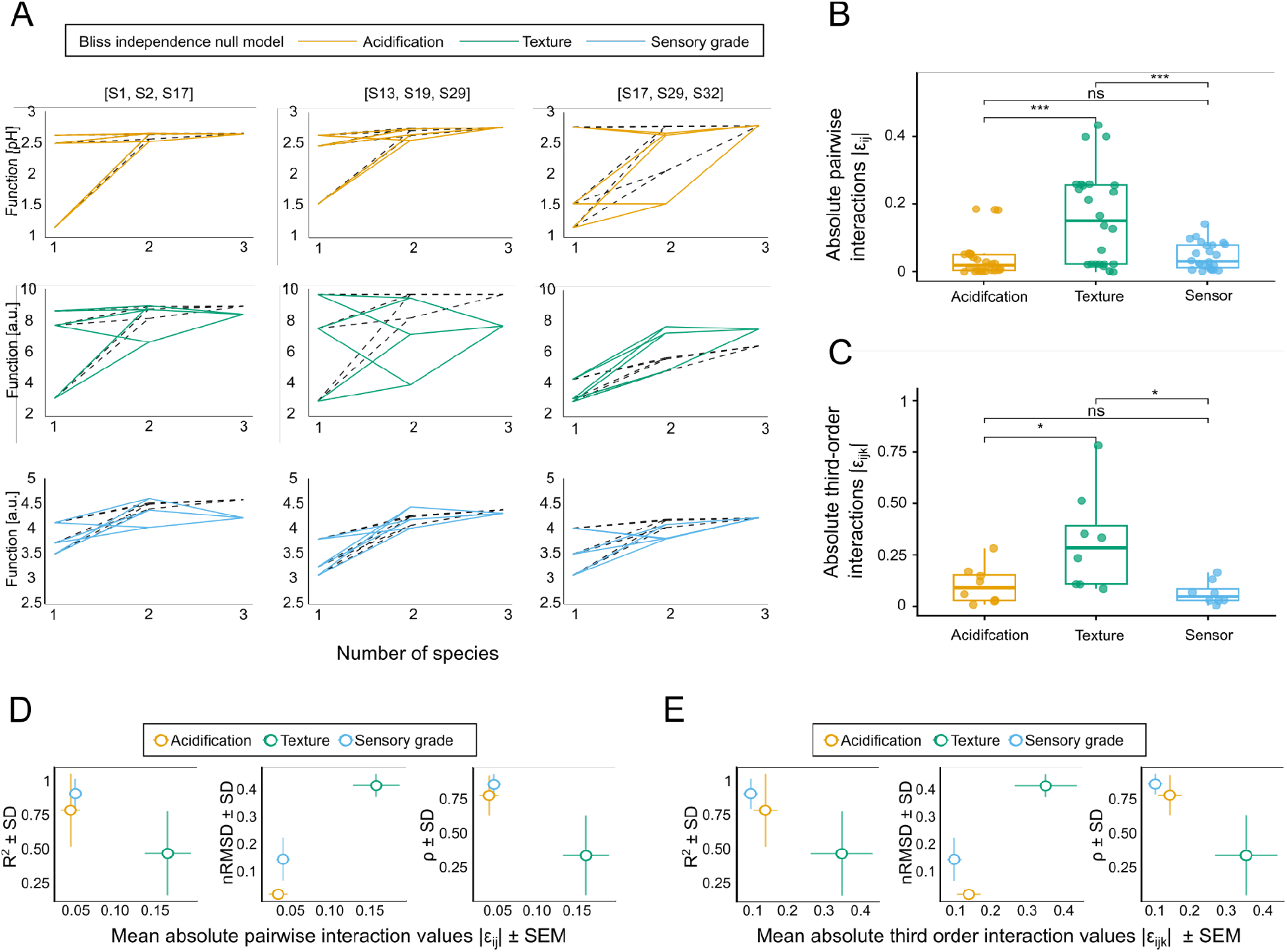
**A)** Three representative fully mapped combinatorial landscapes across three functions. Dashed black lines represent the functional values predicted from the Bliss additive null model, while coloured lines represent the measured function for acidification (orange), texture (green) and sensor grade (blue). Deviations from the null model indicate pairwise and high order interactions shaping the function. **B)** Quantification of pairwise interaction coefficients. Interaction coefficients are quantified as deviations from the Bliss null model across all community compositions for each of the eight subsets. We represented the coefficients as absolute values so that we could compare interaction strengths. Texture (green) exhibits significantly larger pairwise interaction terms compared to acidification (orange) and sensor grade (blue) (pairwise t-test, p<0.01). **C)** Quantification of third order interaction coefficients. Third-order interactions are defined as the deviation of the observed three-member community function from the value expected based on pairwise interactions among its constituent members. Texture shows significantly higher third-order coefficient terms compared to acidification and sensor grade (pairwise t-test, p<0.01). **D)** Relationship between mean absolute pairwise interaction strength and predictive performance for the validation set. The x-axis shows the mean absolute pairwise interaction coefficient, while the y-axis shows predictive performance estimated by bootstrapping the validation set (n=17, 10000 iterations), quantified as R^2^, nRMSD, and Spearman’s ρ. Horizontal error bars indicate the standard error of the mean (SEM) of the absolute pairwise interaction coefficients, and vertical error bars indicate the standard deviation (SD) of the bootstrapped performance estimates. Acidification (orange) and sensory grade (blue) clustered at lower interaction strengths and showed better predictive performance reflected by higher R^2^ and Spearman’s ρ and lower nRMSD. In contrast, texture (green) exhibited higher interaction coefficients and poorer predictive performance relative to the former two functions, with lower R^2^ and Spearman’s ρ and higher nRMSD.**E)**Relationship between mean absolute third order interaction strength and predictive performance for the validation set. The x-axis shows the mean absolute third order interaction coefficient, while the y-axis shows predictive performance estimated by bootstrapping the validation set (n=17, 10000 iterations), quantified as R^2^, nRMSD, and Spearman’s ρ. Horizontal error bars indicate the standard error of the mean (SEM) of the absolute third order interaction coefficients, and vertical error bars indicate the standard deviation (SD) of the bootstrapped performance estimates. Acidification (orange) and sensory grade (blue) clustered at lower interaction strengths and showed better predictive performance reflected by higher R^2^ and Spearman’s ρ and lower nRMSD. In contrast, texture (green) exhibited higher interaction coefficients and poorer predictive performance relative to the former two functions, with lower R^2^ and Spearman’s ρ and higher nRMSD.

**Figure 5.**
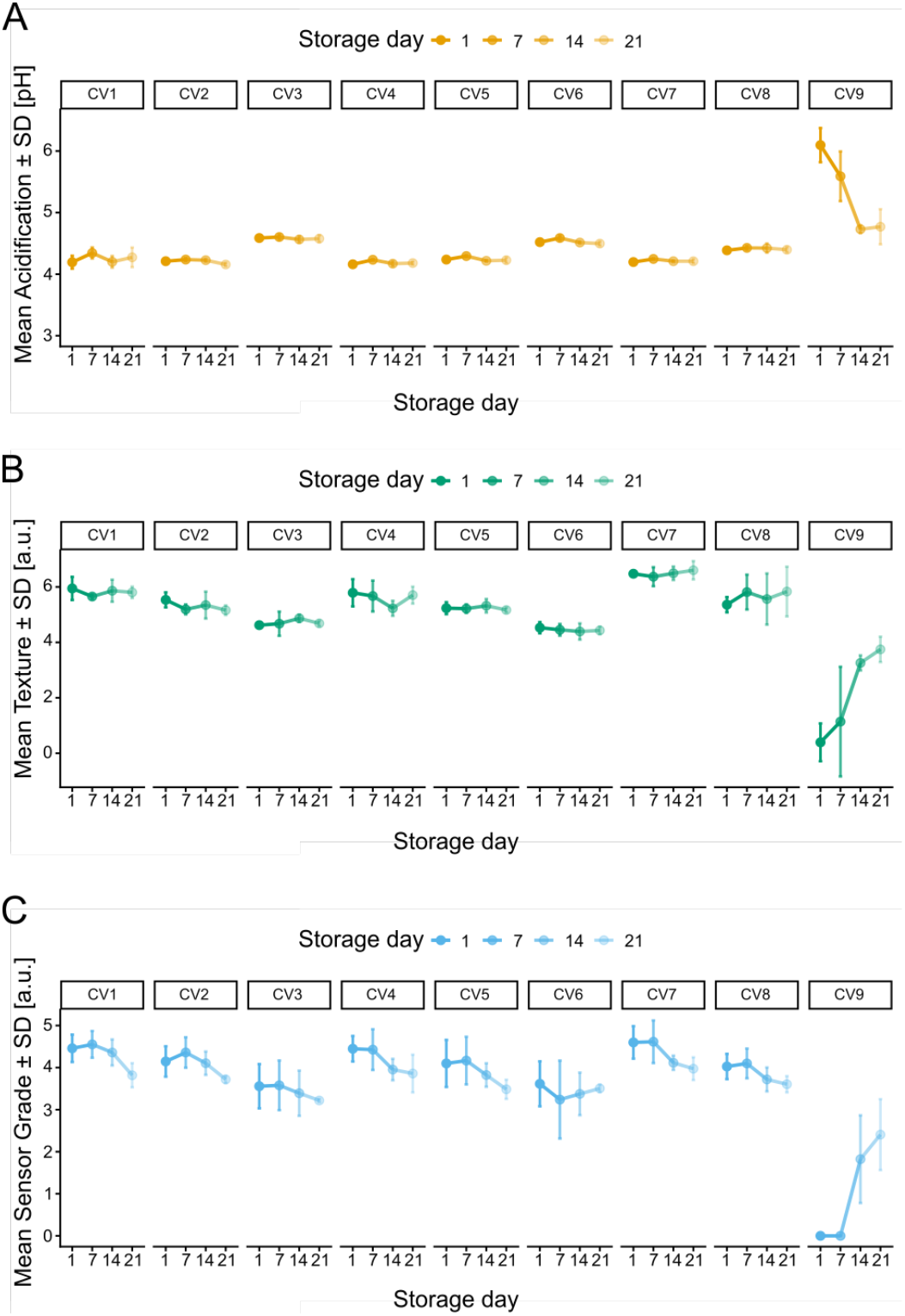
Temporal stability of functional profiles across a 21-day storage time. (A-C) Functional outputs were measured on days 1, 7, 14 and 21 (mean ±SD). Most measured SynCom combinations (8 out of 9) showed relatively stable functional profiles over time, with only modest changes in measured functions. The exception was CV9, characterized by low performance across all measured functions, which exhibited the greatest change during storage, likely due to delayed fermentation kinetics.

Landscape ruggedness did not explain predictability in out-of-sample communities, likely because it additionally depends on the quality of FEE phylogenetic imputation and the degree of conservation (Fig. S10), However, the results were broadly consistent with our hypothesis in the validation communities, with larger average interaction magnitudes (both pairwise and three-order) leading to lower predictability values (Fig. 4C-D). Thus, these results empirically validate previous theoretical results suggesting that more rugged landscapes are inherently less predictable.

### Designed communities retain their functional profiles over time

For practical applications in food fermentation, an important requirement is that designed microbial communities maintain their functional properties over time. We therefore assessed the temporal stability using a subset of nine of the validation communities, over a 21-day storage period. Across all measured functions, functional profiles were broadly stable over time, with all functions showing only modest temporal variation. Notably, the rank ordering of communities was largely preserved (Spearman’sρ= 0.83, 0.87, and 0.93, for acidification, texture, and sensory grade, respectively; p < 0.001 in all cases, n=9), indicating that functional differences established at the time of design persist over time. An apparent exception to the generally observed stability was observed in one low-performing community (CV9), which exhibited stronger temporal changes. However, this change reflected an improvement in function over time rather than degradation, suggesting slower fermentation dynamics as the likely explanation. Overall, functional stability supports the practical applicability of our predictive community design approach.

## Discussion

How accurately can the function of a microbial community be predicted from minimal information about its members? Using soymilk fermentation as a model system with a large biobank of lactic acid bacteria (*53*), we have shown that emergent Functional Effect Equations govern three industrially relevant phenotypes — a metabolic, a physical, and an integrated organoleptic readout — and can be inferred from sparse sampling. Crucially, the slopes and intercepts that parameterize these FEEs are largely phylogenetically conserved across the biobank, allowing us to predict the function of communities composed of previously untested strains from their 16S sequence identity alone. These results position FEE parameters as conserved, species-level ecological traits that can be used for community engineering on a scale at which neither data- and experiment-hungry mechanistic models (*34–37*), machine learning strategies (*44–46*), nor combinatorial screens (*39–42*) are currently tractable. The hypothesis that FEE parameters might themselves be phylogenetically conserved was raised previously in a wine-yeast system (*49*), where despite the conservation of individual functional traits, the conservation of FEE parameters could not be resolved. With FEEs fit on 33 phylogenetically broad strains and validated by imputation onto 17 additional strains drawn from the biobank, our analysis demonstrates that FEE-parameter conservation supports prediction across three industrially relevant community functions.

The accuracy of our predictions varied across the three functions in ways that mapped onto the structure of the underlying landscapes. Acidification was the best-predicted phenotype, and showed landscapes with no large interactions. Texture showed the lowest predictability values and accordingly, its landscapes exhibit substantially stronger pairwise and third-order interactions (Fig. 4B). These results empirically support the hypothesis that landscape ruggedness scales with the biochemical complexity of the function (*55*): acidification reflects a biochemically simple output, while texture depends on a compound set of microbial outputs (e.g. exopolysaccharide production, protease secretion, other secreted metabolites) that might combine in complex ways. Sensory grade, by contrast, seemed to organize on a relatively smooth landscape and was well predicted in the validation communities, which is surprising given that this function might be mechanistically the most complex of all. Moreover, bliss-derived interaction magnitudes correlated with predictive accuracy in within-training validation communities (Fig. 4C-D) but not in out-of-collection communities (Fig. S10), whose accuracy depends additionally on phylogenetic imputation quality. Predictability via conserved FEEs is therefore determined by two factors — landscape ruggedness, which scales with function complexity, and, beyond the training set, degree of conservation and imputation accuracy.

Several limitations of our approach warrant discussion. First, the FEE parameters for each strain were estimated from a moderate number of background communities. While previous work has suggested that this regime is typically sufficient (*48*), some of the strains we identified as phylogenetically discordant might reflect artifacts derived from inaccurate statistical estimation or the influence of outliers (Fig. S5 − S7). Future implementations would benefit from increasing background coverage to disentangle genuine phylogenetic discordance — driven, for instance, by horizontal gene transfer or lineage-specific adaptation — from estimation error. While here we identified these discordant strains post-hoc, future applications in which descriptors are imputed for strains that have not been assayed will require developing confidence measures for imputed parameters, for example based on the phylogenetic distance to the nearest characterized strain. Second, the generalizability of our findings beyond our specific model system and beyond the three functions tested here is ultimately an empirical question. Although global epistasis-like patterns have been reported across diverse systems, the degree to which FEE parameters are phylogenetically conserved may vary substantially across functions and applications. The only previous instance in which this was attempted, using yeast sugar consumption, was inconclusive, and we expect that the parameters for some functions may turn out not to be conserved at all — a possibility consistent with the well-documented heterogeneity of phylogenetic signal in microbial traits more generally (*30,58*). Establishing the generality of the approach will therefore require system-specific testing. Because our out-of-sample validation necessarily probes prediction only within the phylogenetic breadth of the biobank, training collections should be assembled, as ours was, to span the full diversity of the target biobank. Finally, the phylogenetic imputation assumes a Brownian-motion model of trait evolution that may not always hold, and alternative approaches may be considered.

The origin of the conserved scaling between the functional contribution of a species and the function of the community where it is added, described by FEEs, is still not fully understood (*31,48,52*). Theory suggests that these scalings can be understood in terms of effective interactions, which summarize how a species interacts with others at the functional level. Mechanistically, microbial interactions are mediated by traits that are conserved to different degrees(*30,58,60*). Many community functions depend on compound sets of traits that might differ in their degree of conservation. Thus, the question whether emergent traits that summarize species functional effects within communities are themselves conserved remained so far open. Their conservation, which we showed here, opens several avenues of research. First, the heritability of FEE parameters suggests that they behave as bona fide ecological traits, amenable to comparative-genomics approaches (e.g. GWAS-style approaches) that can map their values onto genomic features. Identifying the genomic determinants of FEE slopes and intercepts opens the prospect of elucidating their mechanistic basis, as well as of engineering strains with desired interaction profiles and robust functional outcomes across communities. A second direction concerns the environmental dependence of these descriptors. Because microbial interactions (and their functional outcomes) are sensitive to both biotic and abiotic context (*61,62*), and our FEE parameters were estimated under a single set of fermentation conditions, mapping how these parameters vary across conditions (e.g. temperature, media composition) would delineate to what extent conserved descriptors can be translated between settings (*63*).

The rational design of SynComs underpins a growing range of biotechnological, agricultural, and environmental applications. However, finding which particular combination of strains will deliver the desired function has been limited by a combinatorial gap: modern strain collections vastly exceed the number of consortia that can be experimentally assembled. By describing a species’ functional contribution across communities into two phylogenetically conserved parameters, our findings enable navigating these large community-function landscapes *in silico*, qualitatively changing the economics of SynCom design. More fundamentally, the fact that species’ functional contribution is summarized by heritable quantitative traits has profound implications for our understanding of the evolutionary stability of ecological interactions. Our findings invite a research program in which collective community function can be thought about in the same evolutionary terms as any other organismal trait.

## Materials and methods

### Microorganisms and Fermentation

Thirty-three lactic acid bacteria strains were selected from the LABbank collection of the Institute of Molecular Genetics and Genetic Engineering (*53*) to span the broadest possible phylogenetic diversity within the collection and to include isolates from diverse origin (dairy, silage, plant-derived). The selected panel covered nine genera: *Streptococcus, Lacticaseibacillus, Lactiplantibacillus, Lactobacillus, Lactococcus, Leuconostoc, Limosilactobacillus, Liquorilactobacillus*, and *Enterococcus*. Strains were stored at−80C in MRS or M17 broth with 17% (v/v) glycerol] and propagated in MRS or M17 broth at their species-appropriate optimal growth temperature (30 °C or 37 °C) prior to fermentation experiments.

For the molecular characterization of the isolates, the 16S ribosomal RNA gene was amplified via PCR using specific in-house developed primers UNI16SF (5′-GAGAGTTTGATCCTGGC-3′) and UNI16SR (5′-AGGAGGTGATCCAGCCG-3′). These primers represent shortened, optimized variants of the well-known universal primers 27F and 1540R (*64*). Following amplification, the purified PCR products were sent to Macrogen Inc. (Amsterdam, Netherlands) for Sanger sequencing. To ensure maximum sequence coverage and accuracy, Macrogen utilized the internal sequencing primers 785F and 907R. The generated electropherograms were edited and aligned to construct consensus sequences, which were subsequently analyzed using the BLAST algorithm against the NCBI database.

A complete list of strains, with name, taxonomy, and 16S sequences, is provided in Supplementary Table 3.

### Synthetic community design

For the training experiment, we sampled 307 consortia from the combinatorial space spanned by the 33-strain panel (total 2^33^≈ 10^10^ combinations), including all 33 monocultures and combinations of 2 to 5 members. Community compositions were selected by firstly choosing 9 “background” communities, then assaying all of these with and without the addition of each of the remaining strains. The background communities were chosen to span a range of initial richness (2-4 species), with 4 backgrounds containing 1 species, 3 backgrounds containing 2, 2 backgrounds containing 4. The design made sure that each strain appeared in at least nine independent community background contexts in order to support estimation of strain FEE parameters (*48*) while keeping experimental effort tractable.

For the validation reported in Section 2 of the Results, 17 additional consortia composed of strains from the original 33-strain panel, but not included in any of the 307 training fermentations, were assembled to span the predicted range of community function across all three readouts. To assemble the validation set, microbial consortia were chosen to encompass a full spectrum of performance: high (upper 33%), intermediate (middle 33%), and low (lower 33%) in all three assessed functions (acidification, texture, and sensory grade). The validation set comprised five two-member consortia, six three-member consortia, and six four-member consortia.

For the phylogenetic prediction in out-of-sample SynCosms, 17 additional strains were selected from LABbank that had not been included in any of the 307 training consortia. Strains were selected based on taxonomic congruence with the training set and sequence completeness . Specifically, all 17 isolates are Lactic Acid Bacteria (LAB) belonging to the same genera as the initial 33 strains, and each possesses a near full-length (∼1500 bp) gene sequence to ensure accurate phylogenetic placement. These strains were assembled into 14 validation communities, each containing at least one previously unobserved strain, and 8 of them composed entirely of strains outside of the training set of 33 strains.

For the temporal stability analysis, nine representative SynComs from the validation set of 14 (CV1-CV9) were assayed at four post-fermentation time points (detailed below).

All used SynComs are detailed in Supplementary Table 2.

### Soymilk fermentation

All fermentations were performed in commercial soymilk (Bio Soya Drink, The Bridge S.R.L., Italy; composition: 2.3% fat, 1.4% carbohydrates, 3.8% protein). For each fermentation, overnight cultures in MRS/M17 were inoculated at 3% (v/v) into 15 mL of soymilk in sterile centrifuge tubes. For multi-strain consortia, equal volumes of overnight cultures of each member were combined immediately prior to inoculation resulting in a total 3% (v/v) inoculum. Fermentations proceeded at 42 °C for 15 h without agitation. Each strain combination within the training set *(n* = 307) was prepared in four sterile tubes: one allocated for pH measurement, and the remaining three for textural and sensory analyses. For the validation set and phylogenetic conservation experiment *(n* = 17, *n* = 14), each combination was tested in six sterile tubes, equally divided between pH measurements (three tubes) and textural/sensory assessments (three tubes).

### Functional measurements

#### Acidification

The pH of each fermented sample was measured in triplicate immediately upon completion of fermentation using a Testo 206 pH meter suitable for semi-solid samples (Testo SE & Co. KGaA, Germany). Acidification was quantified as the difference between the pH values of the unfermented soymilk and the post-fermentation pH after 15 hours. This variable was used throughout downstream analyses.

#### Texture (gel hardness)

Texture was assessed in triplicate at room temperature using a TA.XT Plus Texture Analyzer (Stable MicroSystems Ltd., UK) fitted with a 5 kg load cell and a cylindrical penetration probe (P/5, 5 mm diameter). Hardness was defined as the maximum force (in grams) required to drive the probe to a depth of 10 mm at a constant test speed of 1 mm s^−2^, and was extracted using EXPONENT software (Stable MicroSystems Ltd., UK).

#### Sensory grade

Sensory evaluation was performed by a five-member trained panel (two female, three male, aged 25–47) drawn from the Faculty of Agriculture and the Institute of Molecular Genetics and Genetic Engineering. Panelists completed six 2 h training sessions over three weeks using commercial and laboratory-prepared fermented soy products, during which the scoring criteria for four sensory attributes (appearance, odor, texture, and taste) were standardized; the full criteria are given in Supplementary Table 3. Samples were evaluated anonymously on a structured 5-point Likert scale (0 = extremely unacceptable; 5 = extremely acceptable) in 0.25 increments. For each sample and each panelist, an overall acceptability score was computed as the arithmetic mean of the four attribute scores; the sensory grade reported for each sample is the mean of these per-panelist overall scores across the full panel.

For training-set experiments, pH, hardness, and sensory grade were measured at the end of fermentation. For the temporal-stability experiments, all three measurements were repeated at days 1, 7, 14, and 21 of refrigerated storage at 4 °C.

### Functional Effect Equations: estimation and community-function prediction

For each strain *i* and each functional readout, the functional effect of adding strain *i* to a background community **s** was defined as

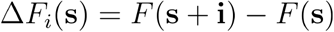

and was regressed against the function of the background community, *F*(**s**) . Functional effect equations (FEEs) were estimated by ordinary least-squares fitting (*48*) of the linear model

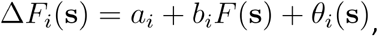

where *a*_*i*_ is the intercept, *b*_*i*_ is the slope, and θ_*i*_(***s***) represent the residuals specific to species *i* in community **s** . Each FEE was fitted using all available (**s**, Δ***F***_***i***_(**s**)) pairs in the training set. The experimental design ensured at least eight independent background contexts per strain (see “Synthetic community design”).

Community function for any consortium was predicted using the “stitching” algorithm introduced in Diaz-Colunga et al. (*48*). Briefly, starting from a measured “seed” community **s**_0_ that is a subset of the target community and whose function has been directly measured, species absent from **s**_0_ are added iteratively. At each step, the function after addition is predicted from the FEE of the added species applied to the function predicted at the previous step. Code implementing FEE estimation and stitching was adapted from the repository (https://github.com/jdiazc9/eco_global_epist) . Predictive accuracy was assessed using three metrics: the coefficient of determination (R^2^), normalized root-mean-square deviation, calculated as

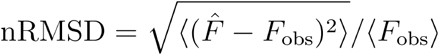

and Spearman correlation. In-sample performance was assessed by 10-fold cross-validation: the 307-consortium training set was randomly partitioned into random 10 folds, with FEEs re-estimated from each 90% partition and used to predict the held-out 10%. This procedure was repeated 500 times in each case. Predictive performance was validated on 17 additional consortia composed of training strains but not present in the training set (Section 2 of Results).

### LASSO regression and prediction

As a benchmark for the FEE-based predictions, we trained a LASSO-regularized linear regression model with explicit pairwise and third-order interaction terms(*55*) on the same 307-consortium training set. Each consortium was encoded as a binary vector x ∈ {–1, +1}^33^ with *x*_*i*_ = +1 denoting presence and *x*_*i*_ = +1 denoting absence of strain *i* (i.e. a Walsh-Fourier expansion of the function in an orthogonal interaction basis). Models were trained for each function using cross-validated LASSO regression, with the regularization parameter selected by the minimum cross-validation error. The fitted models were used to predict the functional output of 17 validation consortia, previously predicted with the FEE-stitching method.

### Phylogenetic tree construction

A maximum-likelihood phylogeny of the 33 LAB strains was constructed from their 16S rRNA gene sequences. Sequences were aligned with MAFFT v7, and algorithm L-INS-i(*65*). The maximum-likelihood tree was inferred with IQ-TREE version 3.1.2 (*66*) using substitution-model selection - GTR. Visualization was performed in R using ggtree (*67*).

### Phylogenetic signal and conservation analysis

Phylogenetic signal in the FEE parameters (slope and intercept) was quantified, for each of the three functions, on the maximum-likelihood 16S phylogeny on the 33-strain panel (see “Phylogenetic tree construction”). Two complementary statistics were used: Abouheif’s *C*_mean_ (*57*), a distance-based index that assumes no explicit model of trait evolution, and Pagel’s λ(*56*), a model-based index derived under Brownian motion. Both were computed with the phylosignal function of the *phylosignal* R package (*68*). For each statistic, significance was assessed by permutation. Trait values were randomized across the tips of the tree (5000 permutations) and the observed value compared against the resulting null distribution, with the p-value defined as the fraction of permutations yielding a statistic at least as extreme as observed. Because six parameters (slope and intercept for three functions) were tested per statistic, p-values were corrected for multiple comparisons within each statistic using the Benjamini-Hochberg method, with significance evaluated at a FDR of 0.05.

To identify strains whose FEE parameters were inconsistent with the surrounding phylogenetic structure, a leave-one-out influence analysis was carried out. Strains were removed one at a time, the signal statistics were recomputed on the remaining 32 strains (999 permutations), and the change relative to the full-panel estimate, Δ = (signal without the focal strain) − (full-panel signal), was recorded for every parameter and statistic. A strain was classified as phylogenetically discordant when its removal produced a marked increase in signal (large positive Δ) concordantly across both statistics and across multiple functions. Three strains (S8, S14 and S22) emerged as consistently and substantially the most influential by this criterion (Supplementary Figure S5 and S6) and were excluded from the conservation analysis.

Phylogenetic signal was then re-estimated on the resulting 30-strain (“pruned”) tree and compared with the full-panel estimates (Fig. 3B).

### Phylogenetic imputation of FEE parameters and out-of-collection validation

To test whether FEE parameters could be predicted for previously unassayed strains, slope and intercept values were imputed for the 17 out-of-sample strains from their positions on the extended 50-taxon 16S phylogeny, comprising the 33-strain panel together with the 17 out-of-sample strains. Prior to imputation the tree was rendered fully bifurcating (multi2di function), zero-length branches were set to 10^−6^, and the tree was made ultrametric by penalized log-likelihood (function chronos in package *ape*(*69*)) assuming that trait divergence accrues with evolutionary time instead of lineage specific 16S substitution accumulation rate. Each FEE parameter was then imputed independently under a univariate Brownian-motion model (phylopars function, *Rphylopars* R package(*70*)), with the parameter values of the out-of-sample strains were treated as missing and predicted as their reconstructed values at the corresponding tips, with the 30 experimentally assayed strains that were retained in the “pruned” tree providing the observed data.

Community function for the 14 validation consortia containing previously unobserved strains was then predicted with the FEE stitching procedure described above, using the imputed slope and intercept values for the unassayed strains; predicted and measured functions were compared and accuracy quantified by R^2^ and nRMSD as defined above.

### Quantification of pairwise and higher-order interactions

To quantify pairwise and third-order functional interactions for each readout, we used eight three-species subsets of the 33-strain panel for which the full combinatorial landscape was experimentally characterized: each subset’s three monocultures, all three two-member combinations, and the three-member community.

For each subset and each readout, observed function values were rescaled to a bounded effect fraction

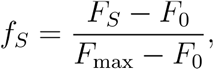

where *F*_*S*_ is the observed function of consortium *S*,*F*_*0*_ = 0 is the empty-community baseline, and *F*_max_ is the maximum function observed across consortia in the subset. We denote the rescaled monoculture function of species *i* by *f*_*i*_.

Under Bliss independence, the expected effect of a consortium *S* with no interactions among its members is

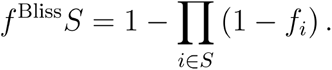

We chose a Bliss null because our functions are bounded. The fraction *f* ∈ [0, 1] is multiplicative under independence, and Bliss is guaranteed to respect this bound.

The net deviation from the null for any consortium was quantified as

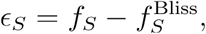

with *ϵ*_*s*_ > 0 indicating synergy and ***ϵ***_***s***_ < 0 antagonism. Pairwise interaction coefficients ***ϵ***_*ij*_ were calculated directly from the two-member consortia. For three-member consortia, we partitioned the net deviation ***ϵ***_*ij*k_ into a pairwise component and a third-order interaction component (*71,72*):

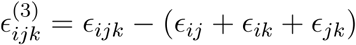

Under this convention, 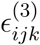 isolates the component of the three-member interaction that cannot be accounted for by single effects nor by pairwise interactions.

Interaction magnitudes used in the comparisons across readouts (Fig. 4B–E) were absolute values of ***ϵ***_*ij*_ and 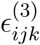, computed separately for each of the three functions.

### Bootstrapping

Due to relatively small sample sizes of the validation and the out-of-sample SynCom sets, comprising 17 and 14 communities, respectively, we used bootstrapping to assess the robustness of the obtained performance estimates. This analysis was performed for the FEE-based predictions in both the validation and out-of-sample set, and for the LASSO predictions of the validation set. For each set and function, communities were resampled with replacement 10 000 times. In each iteration, prediction performance was recalculated from the resampled observed and predicted values. Performance was quantified using three metrics: R^2^, nRMSD, and Spearman’s ρ.

## Supporting information

Suppelementary Material

## Data and code availability

All raw experimental data and analysis code generated in this study are publicly available at [https://github.com/gojkovic10/Soymilk]. Code implementing FEE estimation and stitching was adapted from the eco_global_epist repository (https://github.com/jdiazc9/eco_global_epist) accompanying (*48*). All analyses were performed in R [PLACEHOLDER: version] and the following specific packages: glmnet, phytools, adephylo, ape, vegan, ARTool, ggpubr, Rphylopars, readxl, dplyr, ggplot2, scales, tibble, grid, combinat, MASS, testthat, ggbeeswarm, tidyr, stringr, and patchwork .

The authors used an AI model (Claude Opus 4.7, Anthropic) for coding assistance, to help edit text for readability and conciseness in several sections of the manuscript, and to generate the cartoon images for the three functions in Figure 1A. All scientific content was written, verified, and approved by the authors.

## Author contributions

Conceptualization: Nemanja Stanisavljević, Nemanja Kljajević, Djordje Bajic Data Curation: Uroš Gojković, Nemanja Kljajević, Djordje Bajic

Formal Analysis: Uroš Gojković, Nemanja Stanisavljević, Zorana Miloradović, Nemanja Kljajević, Nikola Popović, Goran Vukotić, Nina Medaković, Djordje Bajic

Funding Acquisition: Nemanja Stanisavljević, Zorana Miloradović, Djordje Bajic

Investigation: Uroš Gojković, Nemanja Stanisavljević, Zorana Miloradović, Nemanja Kljajević, Nikola Popović, Goran Vukotić, Nina Medaković, Djordje Bajic

Methodology: Uroš Gojković, Nemanja Kljajević, Nemanja Stanisavljević, Djordje Bajic Project Administration: Nemanja Stanisavljević, Djordje Bajic

Resources: Zorana Miloradović, Nikola Popović Supervision: Nemanja Stanisavljević, Djordje Bajic Software and visualization: Uroš Gojković, Djordje Bajic

Writing – Original Draft Preparation: Uroš Gojković, Nemanja Kljajević, Nemanja Stanisavljević, Djordje Bajic

Writing – Review & Editing: Uroš Gojković, Nemanja Kljajević, Nemanja Stanisavljević, Goran Vukotić, Djordje Bajic

## Acknowledgements

We thank Juan Diaz-Colunga, members of the Industrial Microbiology Section at TU Delft, Amarela Terzić-Vidojević and members of the Probiotics and Microbiota-Host Interaction group and Molecular Microbiology group at IMGGE, for useful feedback and discussions. NP, NM, NS, and NK were supported by the Ministry of Science, Technological Development, and Innovation of the Republic of Serbia, Grant 451-03-33/2026-03/200042. GV was supported by the Ministry of Science, Technological Development, and Innovation of the Republic of Serbia, Grant No. 451-03-34/2026-03/200178. ZM was supported by the Ministry of Science, Technological Development, and Innovation of the Republic of Serbia, Grant No. 451-03-33/2026-03/200116. DB wants to thank the Zero Emission Biotechnology program from the Department of Biotechnology at TU Delft for support.

## Supplementary Information

**Figure S1.**
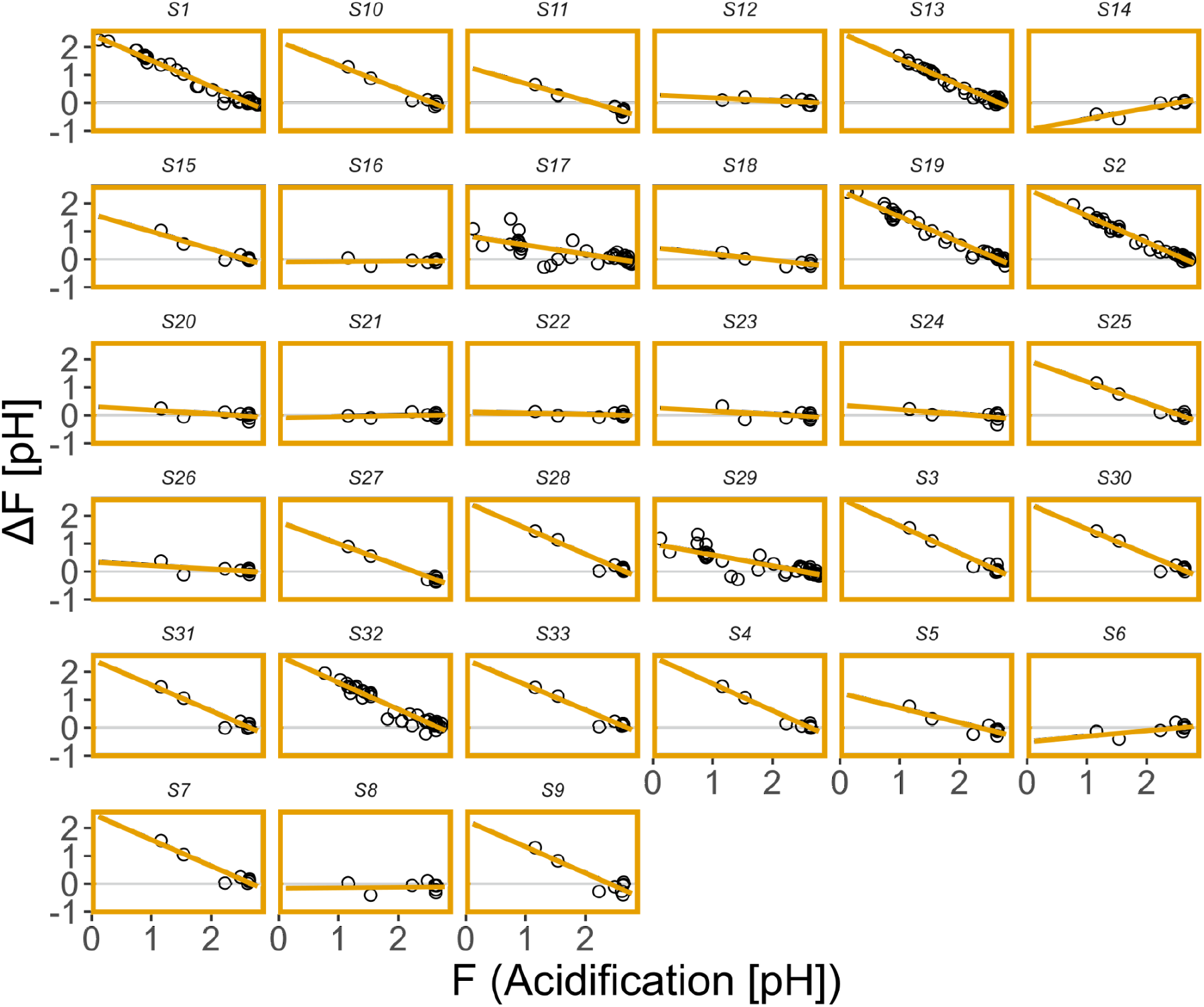
Functional effect equations (FEEs) constructed for acidification. For each of the 33 assayed LAB strains, we have constructed FEEs based on the effect of each of the strains, with each strain assayed on at least 8 community backgrounds. The functional effect (ΔF) is defined as the difference in community function between the background alone and the background with the focal species.

**Figure S2.**
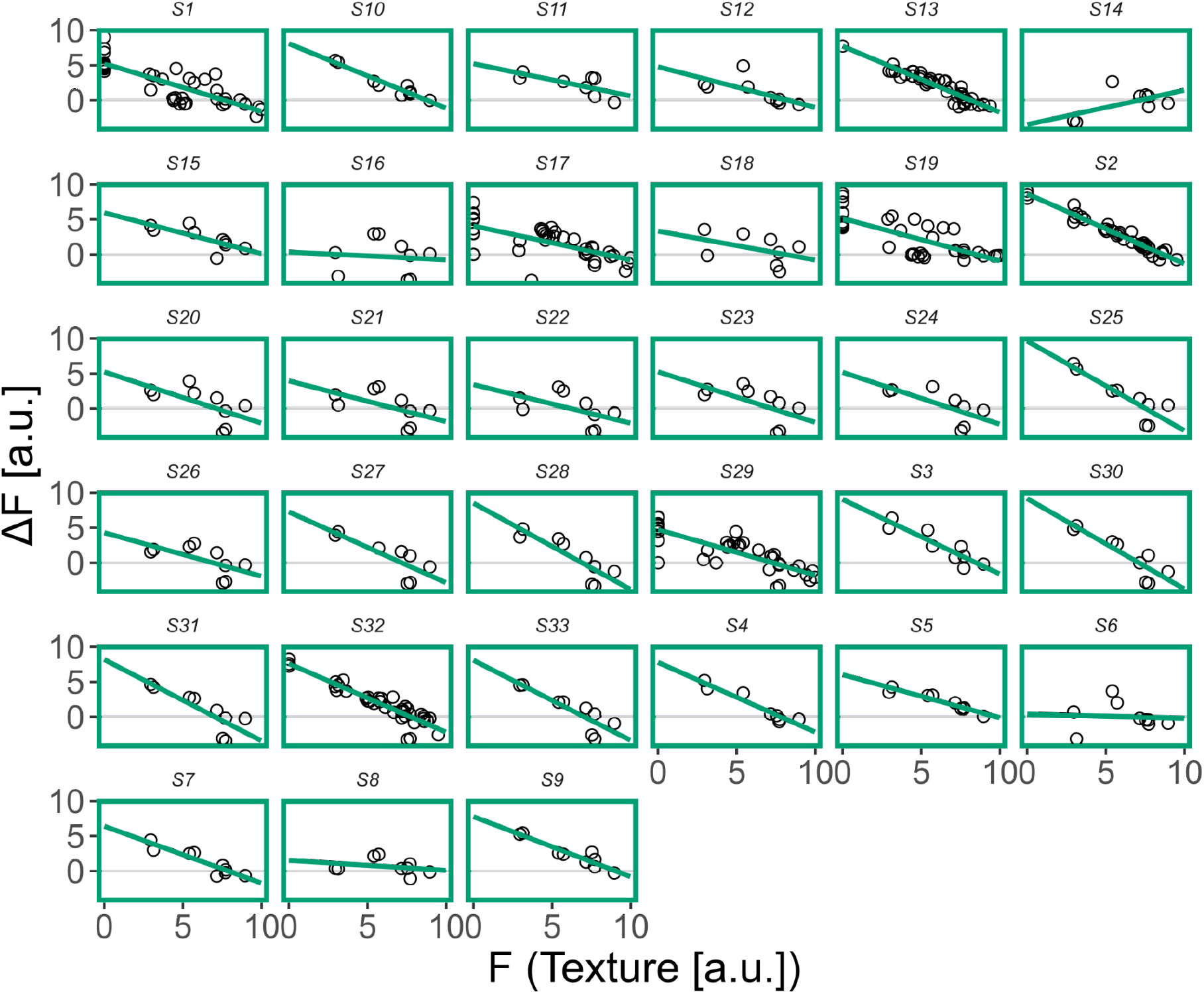
Functional effect equations (FEEs) constructed for texture. For each of the 33 assayed LAB strains, we have constructed FEEs based on the effect of each of the strains, with each strain assayed on at least 8 community backgrounds. The functional effect (ΔF) is defined as the difference in community function between the background alone and the background with the focal species.

**Figure S3.**
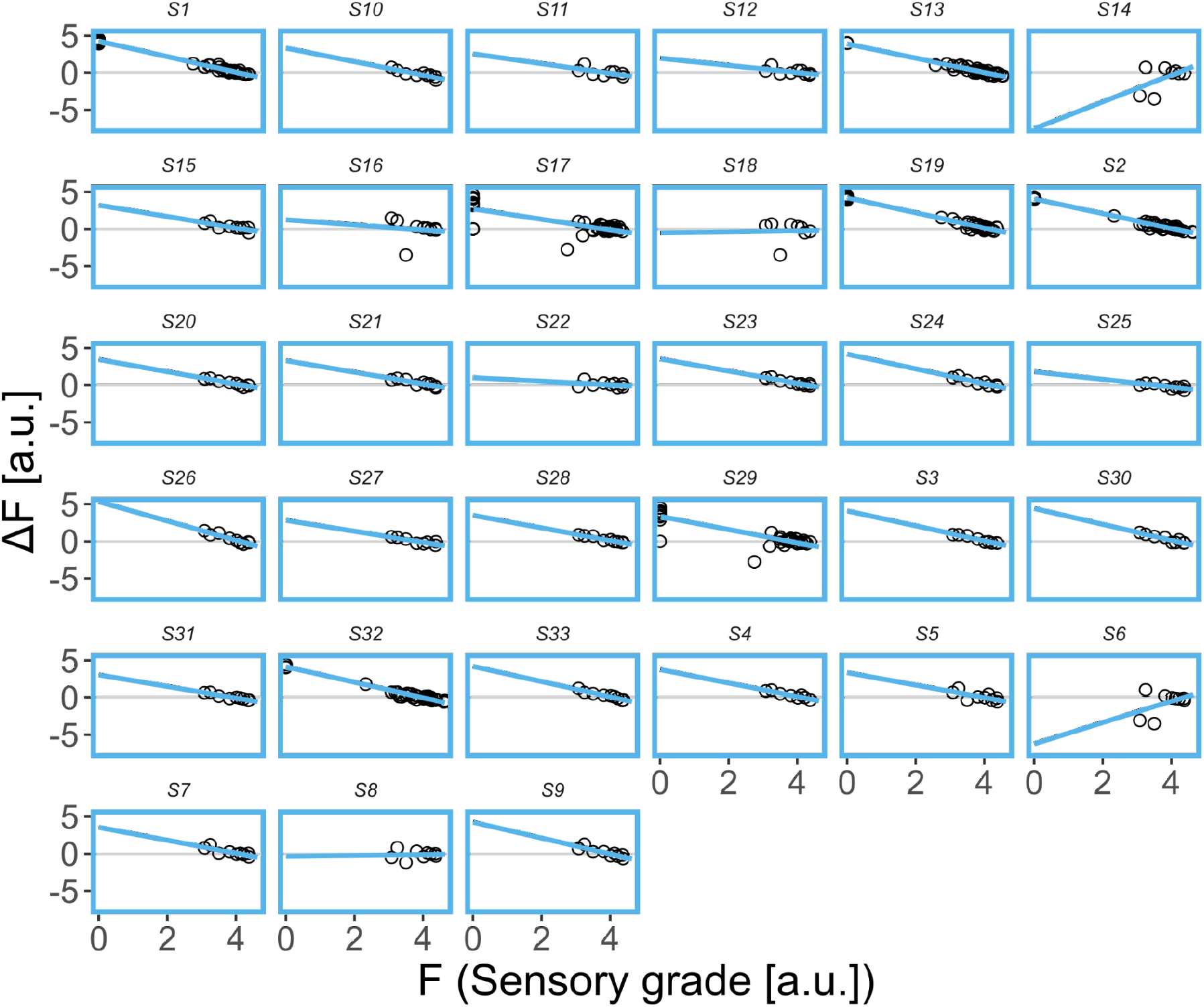
Functional effect equations (FEEs) constructed for sensor grade. For each of the 33 assayed LAB strains, we have constructed FEEs based on the effect of each of the strains, with each strain assayed on at least 8 community backgrounds. The functional effect (ΔF) is defined as the difference in community function between the background alone and the background with the focal species.

**Figure S4.**
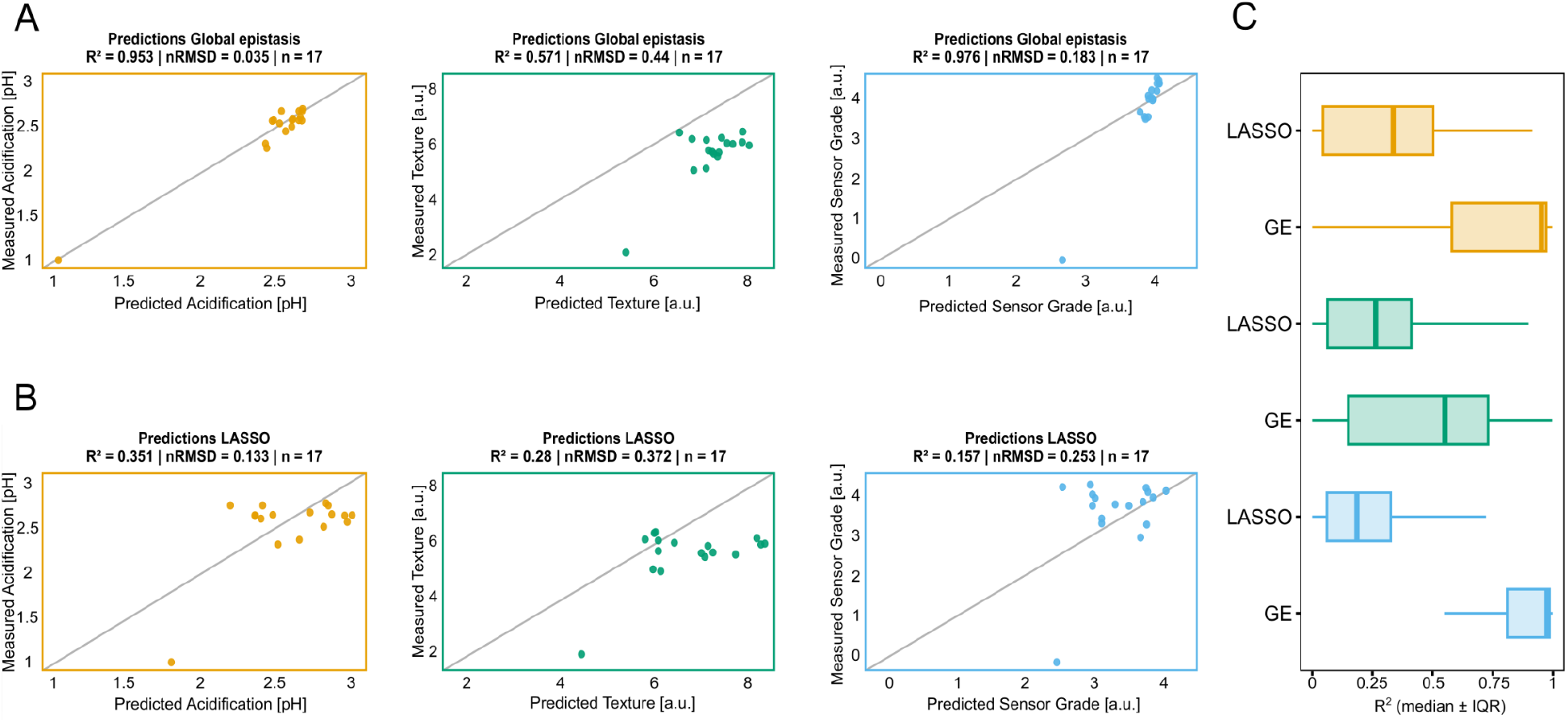
Detailed comparison between FEE-based and LASSO predictions. **A-B)** Predicted versus measured values using FEE stitching (A) or LASSO (B) on the validation set for the three functions (N = 17 in each case). Note that the panels in A reproduce the same coloured points shown in Fig. 2A (main text), represented here separately to allow direct comparison with LASSO predictions (panel B). **C)** Comparison of bootstrapped R^2^ values (10000 iterations) between the FEE-stitching model and LASSO regression. Colors represent function, acidification (orange), texture (green) and sensor grade (blue). FEE-stitching consistently outperforms LASSO for the R^2^ metric in all three functions (t-test p<10^−16^ in all cases).

**Figure S5.**
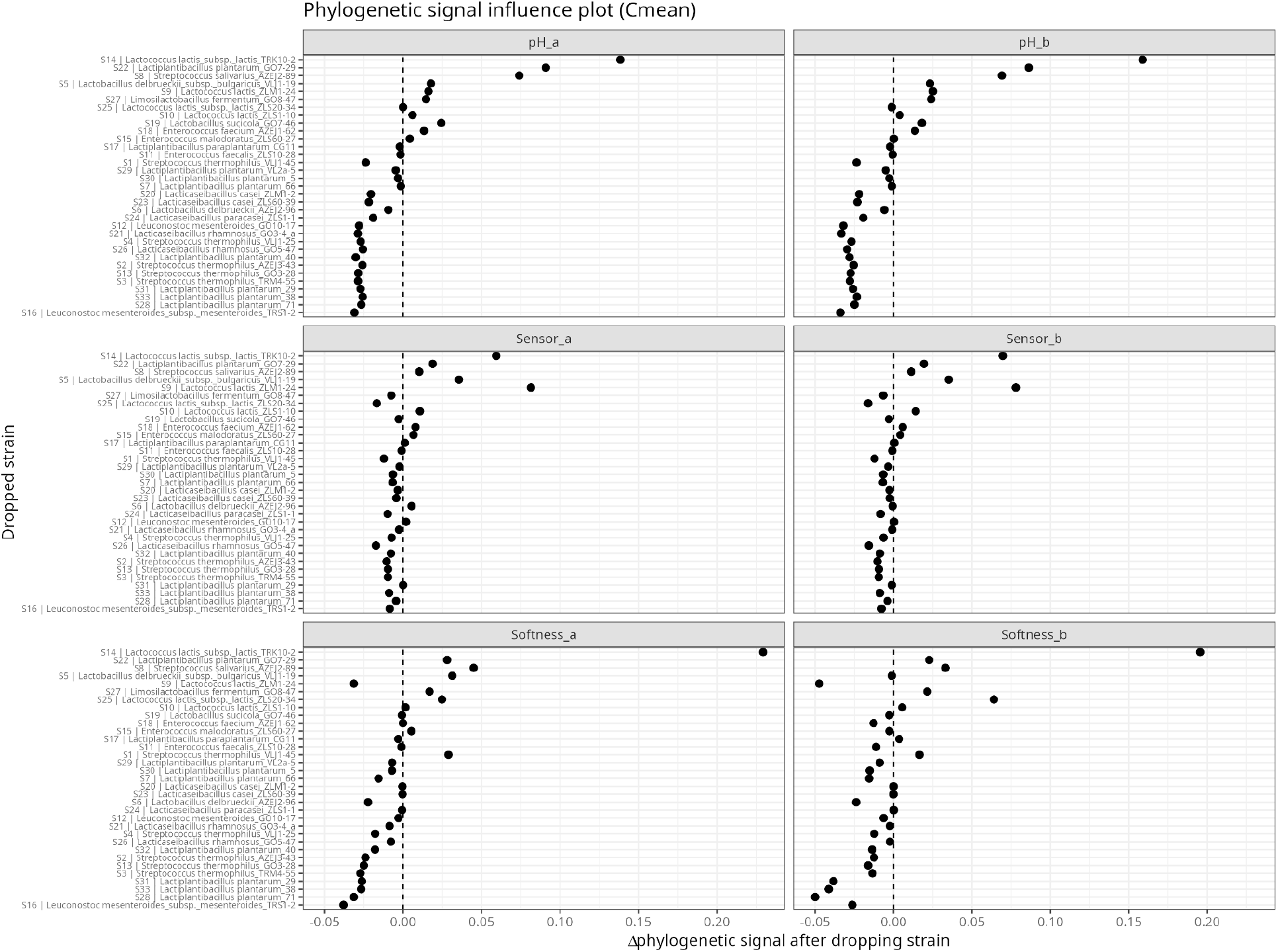
Leave-one-out influence analysis of phylogenetic signal *(C*_mean_) in FEE parameters. Each panel corresponds to one FEE parameter (slope, *a*, or intercept, *b)* for acidification (pH), texture (Softness) and sensory grade (Sensory grade). Points show the change in phylogenetic signal, Δ = (signal estimated after removing the focal strain) − (signal estimated on the full 33-strain panel), with the signal measured as Abouheif’s *C*_mean_. Positive Δ indicates strains whose removal increases the signal (i.e. isolates departing from phylogenetic structure) whereas negative Δ indicates strains that reinforce phylogenetic trait structure. The dashed line marks Δ = 0. Strains are ordered along the y-axis by their mean Δ across all six parameters. Three isolates (S8, S14 and S22) show the largest positive Δ across multiple parameters and functions, identifying them as phylogenetically discordant; these were excluded from the conservation analysis (“pruned” tree, see main text).

**Figure S6.**
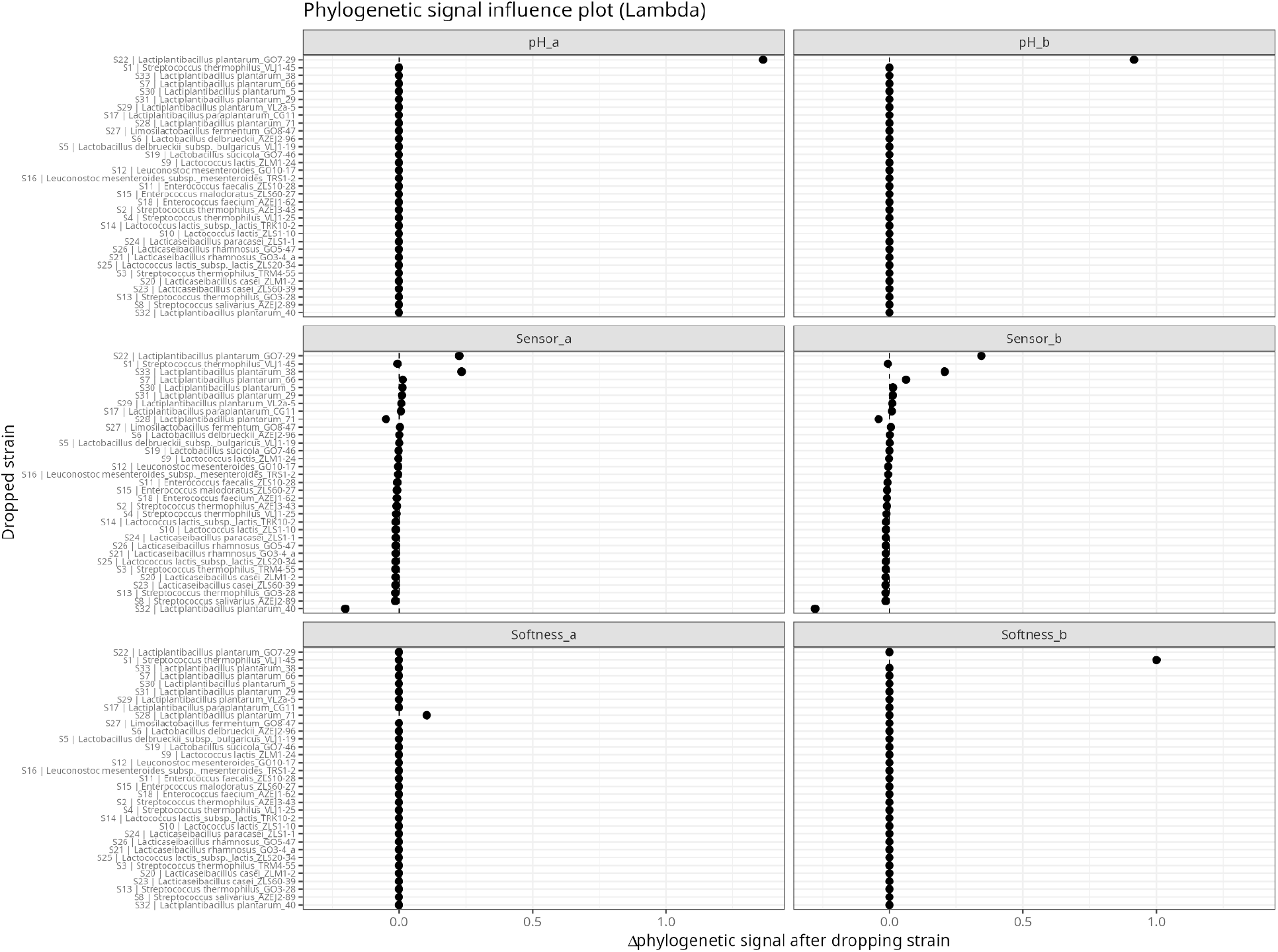
Leave-one-out influence analysis of phylogenetic signal (Pagel’s λ) in FEE parameters. Figure analogous to Fig S5 for the Pagel’s λ indicator.

**Figure S7.**
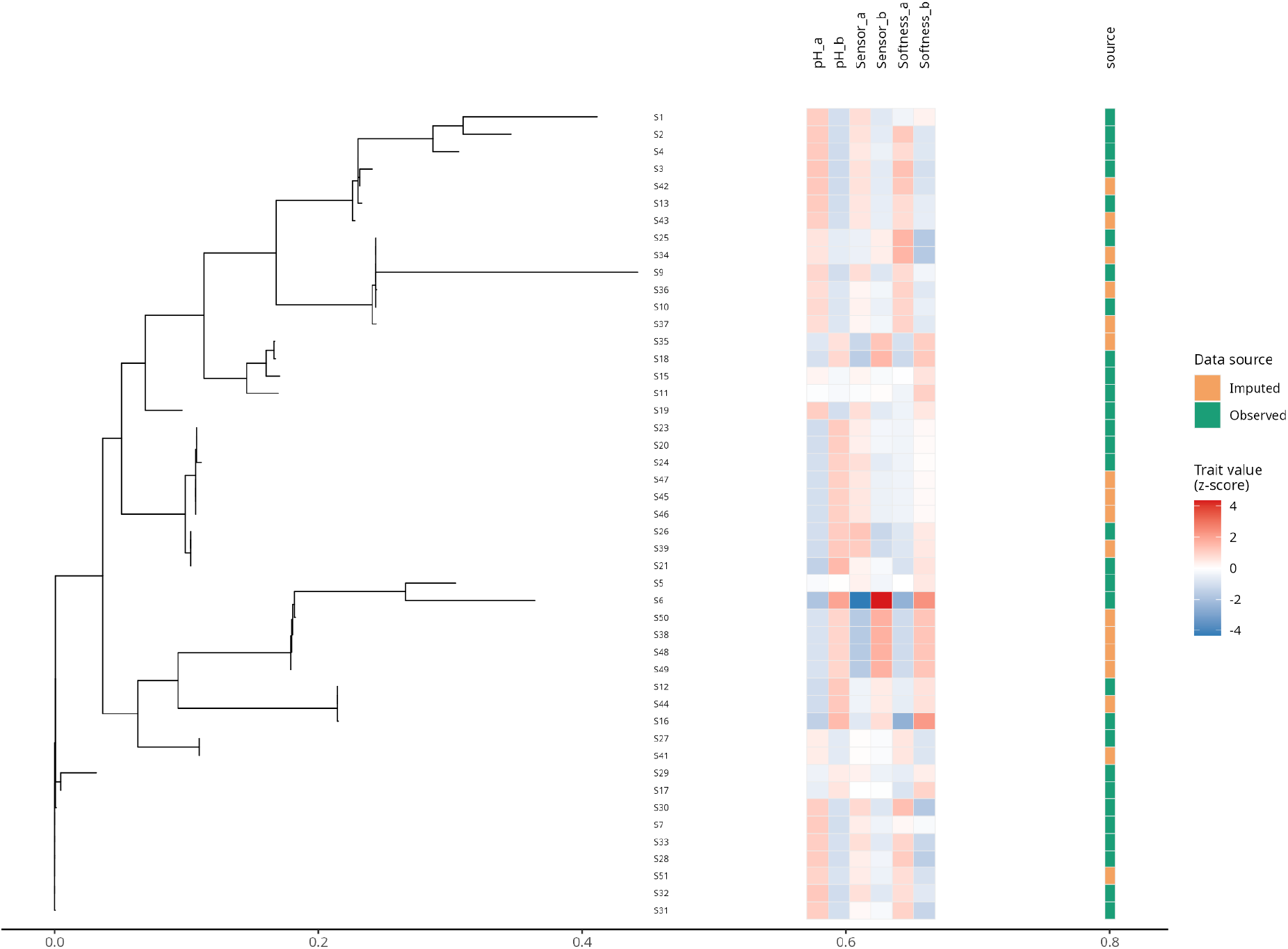
Full phylogeny including imputed trait values for out of sample strains. Phylogenetic tree analogous to Fig. 3A, with strains S8, S14 and S22 removed (“pruned” tree) and including the out-of-sample strains. The “source” column indicates whether a strain belongs to the initial pool (“Observed”) or to the out-of-sample pool (“Imputed”).

**Figure S8.**
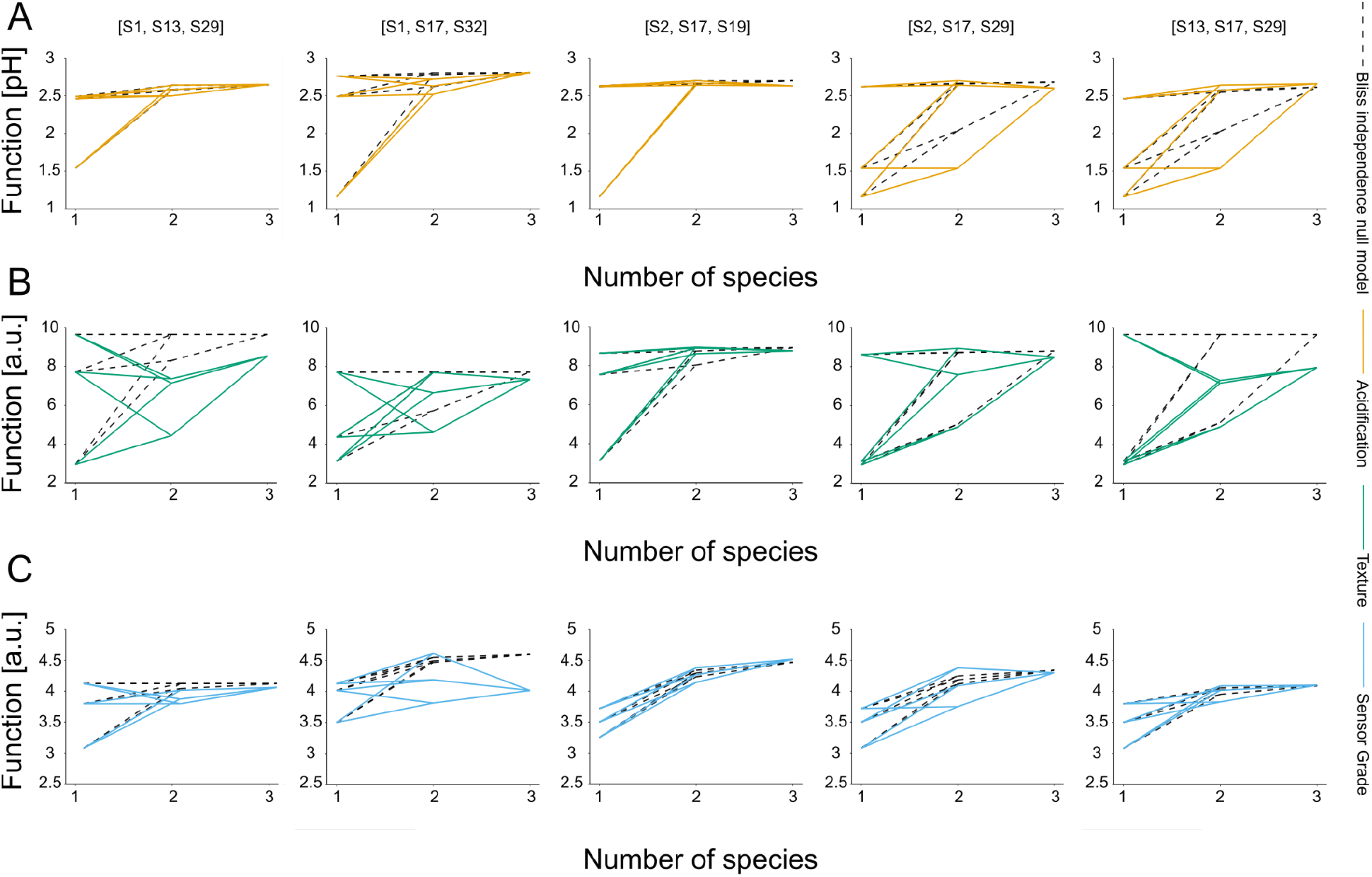
Fully mapped combinatorial landscapes across measured functions. (A-C) Functional landscapes for 5 out 8 subsets used for interaction coefficient calculations (with the remaining 3 shown in Fig. 4A, main text). Dashed black lines represent the predicted values of two- and three-member SynComs based on the Bliss independence null model (Methods), while coloured lines represent the measured functions for acidification (orange), texture (green) and sensor grade (blue). Deviations from the null model define interactions.

**Figure S9.**
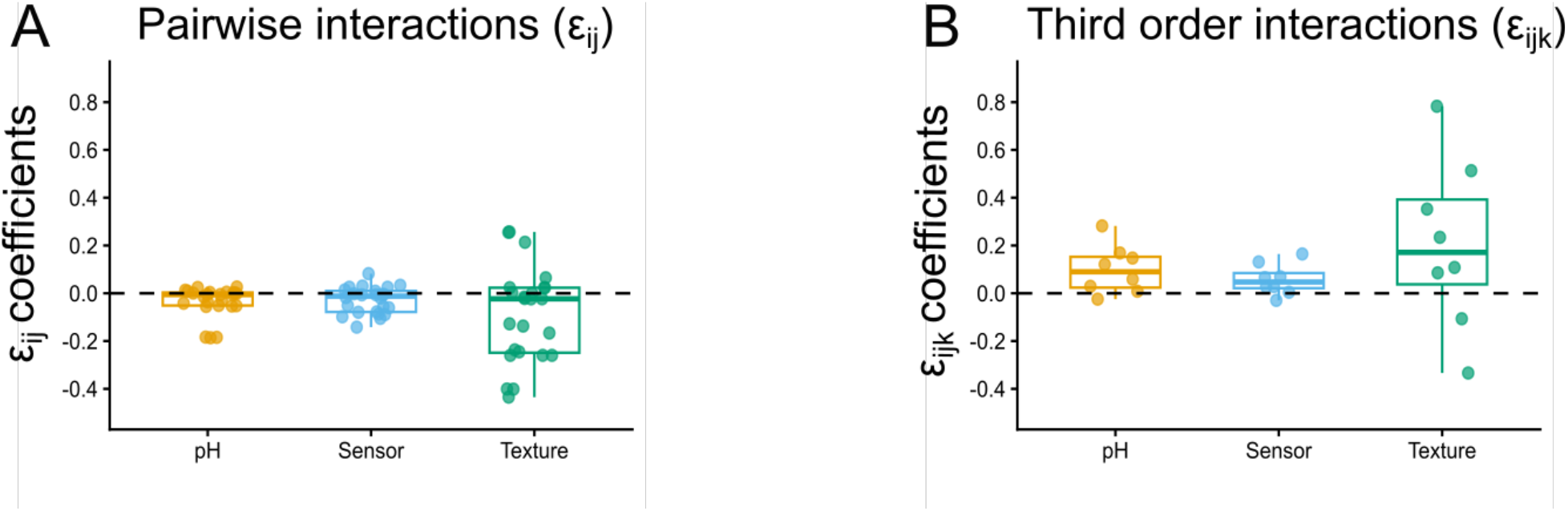
**A)** Non-absolute values of pairwise interaction coefficient, quantified as deviations from the Bliss null model across all community compositions for each of the eight subsets. **B)** Non-absolute values of third order interaction coefficients. Third-order interactions are defined as the deviation of the observed three-member community function from the value expected based on pairwise interactions among its constituent members.

**Figure S10.**
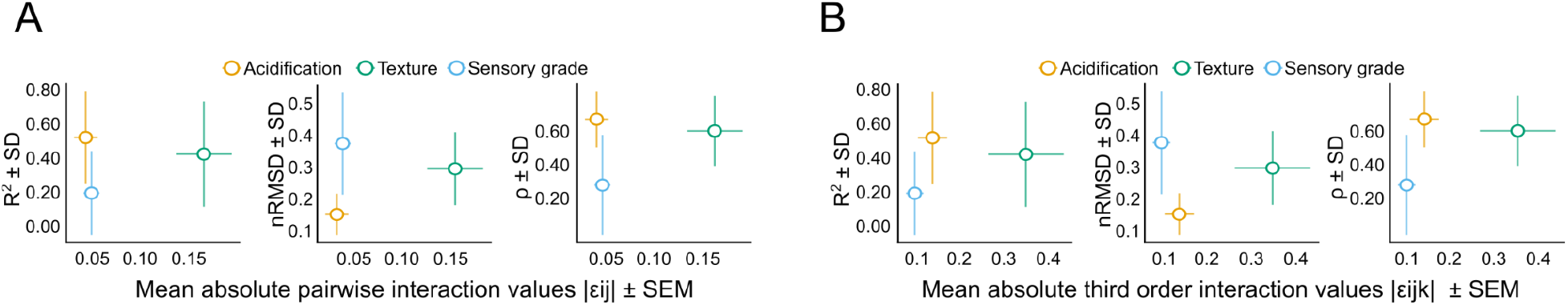
**A)** Relationship between mean absolute pairwise interaction strength and predictive performance for the out-of-sample set of SynComs (i.e. those including phylogenetically imputed FEEs). The x-axis shows the mean absolute pairwise interaction coefficient, while the y-axis shows predictive performance estimated by bootstrapping the out of sample set (N=14, 10000 iterations), quantified as R^2^, nRMSD, and Spearman’s ρ. Horizontal error bars indicate the standard error of the mean (SEM) of the absolute pairwise interaction coefficients, and vertical error bars indicate the standard deviation (SD) of the bootstrapped performance estimates. **B)** Relationship between mean absolute third order interaction strength and predictive performance for the out of sample set. The x-axis shows the mean absolute third order interaction coefficient, while the y-axis shows predictive performance estimated by bootstrapping the out of sample set (N=14, 10000 iterations), quantified as R^2^, nRMSD, and Spearman’s ρ. Horizontal error bars indicate the standard error of the mean (SEM) of the absolute third order interaction coefficients, and vertical error bars indicate the standard deviation (SD) of the bootstrapped performance estimates. In all plots, orange dots represent acidification, green texture and blue sensory grade.

**Supplementary table 1.**
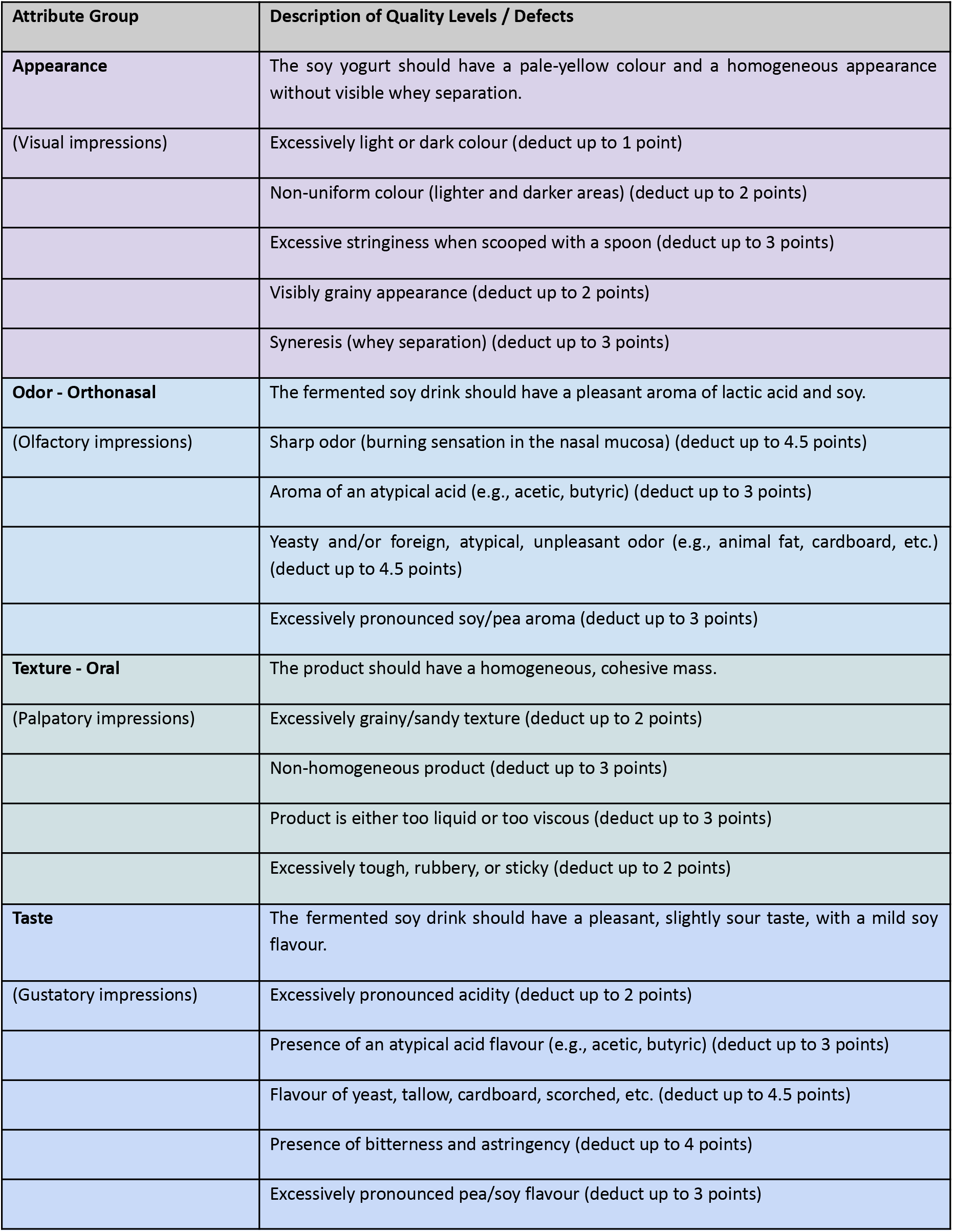
Grading panel for Sensory grade function. The table shows the criteria used to evaluate soy yoghurt quality. Fermented soy milk products were scored based on appearance, odor, oral texture and taste. For each category, desirable product characteristics and specific sensory defects are listed together with the corresponding point deductions applied during scoring.

**Supplementary Table 2.**
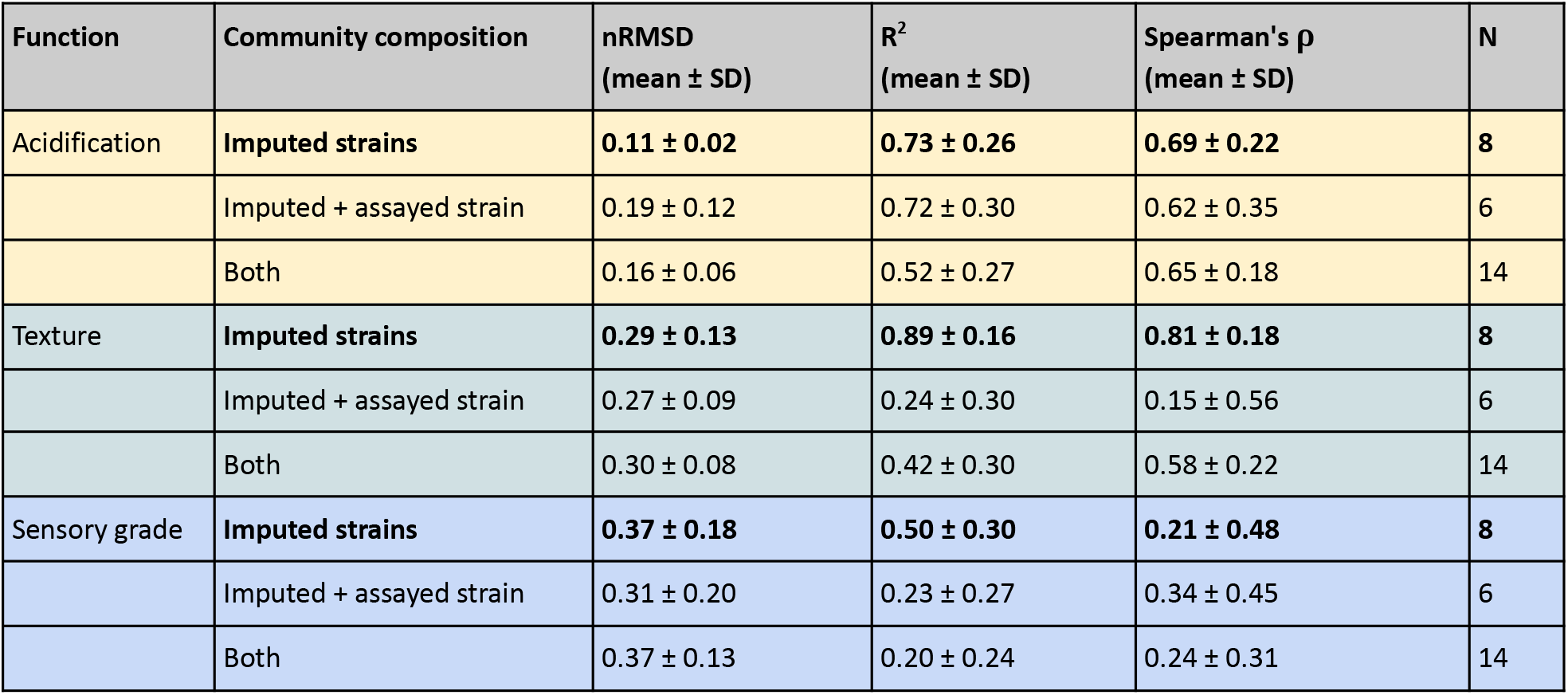
Bootstrapped prediction performance for the out of sample set. The table summarizes prediction performance for the out of sample set containing only imputed strains, communities containing both imputed and assayed strains, and the combined set of both community types. Performance is reported for three measured functions: acidification, texture and sensory grade. For each function and community type, communities were reasampled with replacement for 10000 bootstrap iterations. Mean ± SD values are shown for nRMSD, R^2^, and Spearman’s rank correlation coefficient (ρ). The number of communities in each community type is indicated by N. Bolded rows correspond to values reported in Fig. 3C-E of the main text.

| Function | Community composition | nRMSD<br>(mean ± SD) | R<sup>2</sup><br>(mean ± SD) | Spearman's $\rho$<br>(mean ± SD) | N |
| --- | --- | --- | --- | --- | --- |
| Acidification | **Imputed strains** | **0.11 ± 0.02** | **0.73 ± 0.26** | **0.69 ± 0.22** | **8** |
| | Imputed + assayed strain | 0.19 ± 0.12 | 0.72 ± 0.30 | 0.62 ± 0.35 | 6 |
| | Both | 0.16 ± 0.06 | 0.52 ± 0.27 | 0.65 ± 0.18 | 14 |
| Texture | **Imputed strains** | **0.29 ± 0.13** | **0.89 ± 0.16** | **0.81 ± 0.18** | **8** |
| | Imputed + assayed strain | 0.27 ± 0.09 | 0.24 ± 0.30 | 0.15 ± 0.56 | 6 |
| | Both | 0.30 ± 0.08 | 0.42 ± 0.30 | 0.58 ± 0.22 | 14 |
| Sensory grade | **Imputed strains** | **0.37 ± 0.18** | **0.50 ± 0.30** | **0.21 ± 0.48** | **8** |
| | Imputed + assayed strain | 0.31 ± 0.20 | 0.23 ± 0.27 | 0.34 ± 0.45 | 6 |
| | Both | 0.37 ± 0.13 | 0.20 ± 0.24 | 0.24 ± 0.31 | 14 |

**Supplementary Table 3 (provided as supplementary file in excel: “Supplementary table 3 - strain annotation + 16s sequences”). Strain notation and taxonomy**. The table lists the strain notation IDs used throughout the study, the corresponding taxonomic names, and the associated 16S rRNA gene sequences. Strains denoted S1-S33 refer to the assayed strains, whereas S34-S51 refer to strains with phylogenetically imputed FEEs.

**Supplementary Table 4 (provided as** supplementary **file in excel: “Supplementary table 4 - Community annotation”). Strain composition of tested communities**. The table lists the community notation IDs used throughout the study, the corresponding strain composition, and the community type. Community notation IDs starting with C refer to communities used in the training dataset (N = 307), IDs beginning with CV refer to validation set communities (N=17) and IDs beginning with COS refer to out-of-sample communities (i.e. those including strains with phylogenetically imputed FEEs, N=14).

