## Supplementary material for "Conserved emergent traits enable phylogenetic prediction of community function": Suppelementary Material

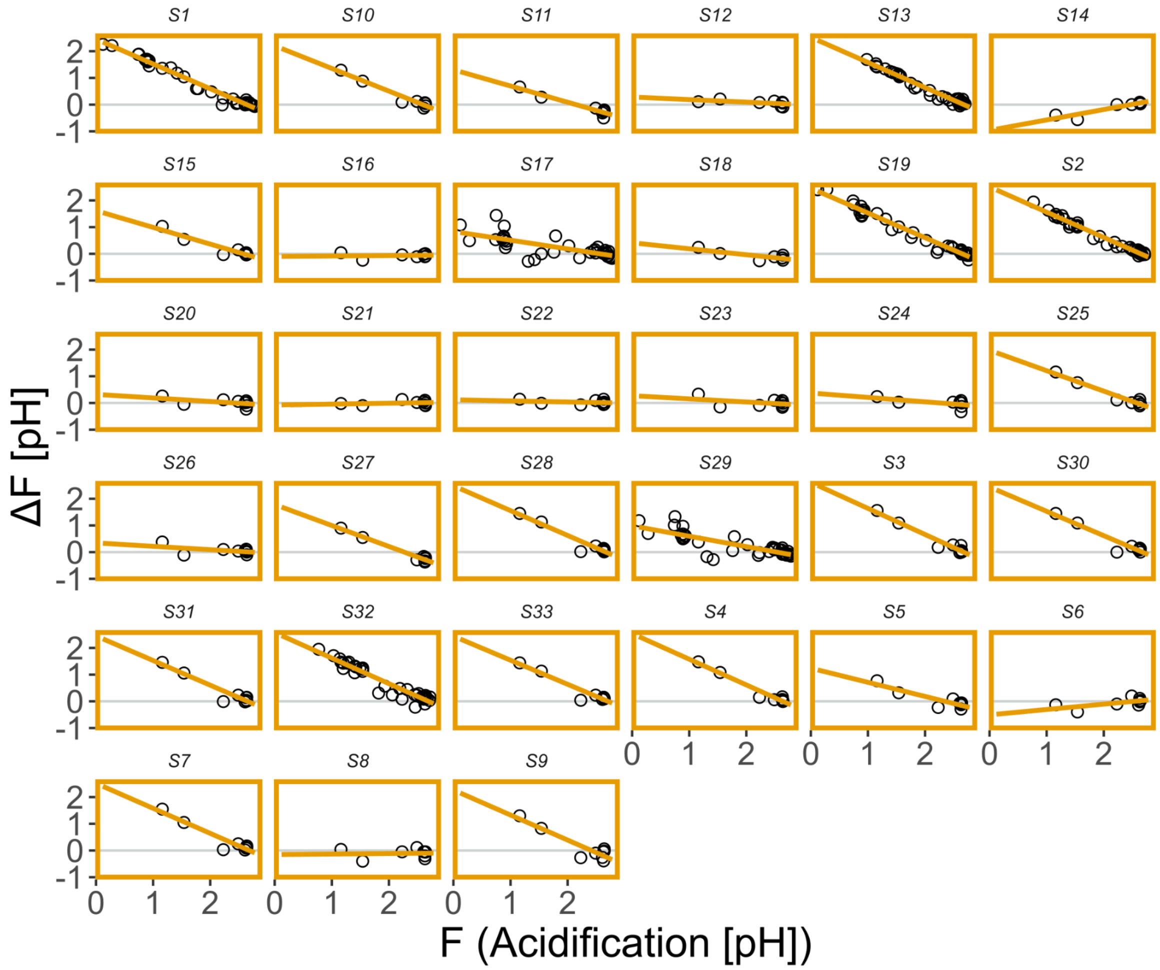


**Figure S1. Functional effect equations (FEEs) constructed for acidification.** For each of the 33 assayed LAB strains, we have constructed FEEs based on the effect of each of the strains, with each strain assayed on at least 8 community backgrounds. The functional effect (ΔF) is defined as the difference in community function between the background alone and the background with the focal species.

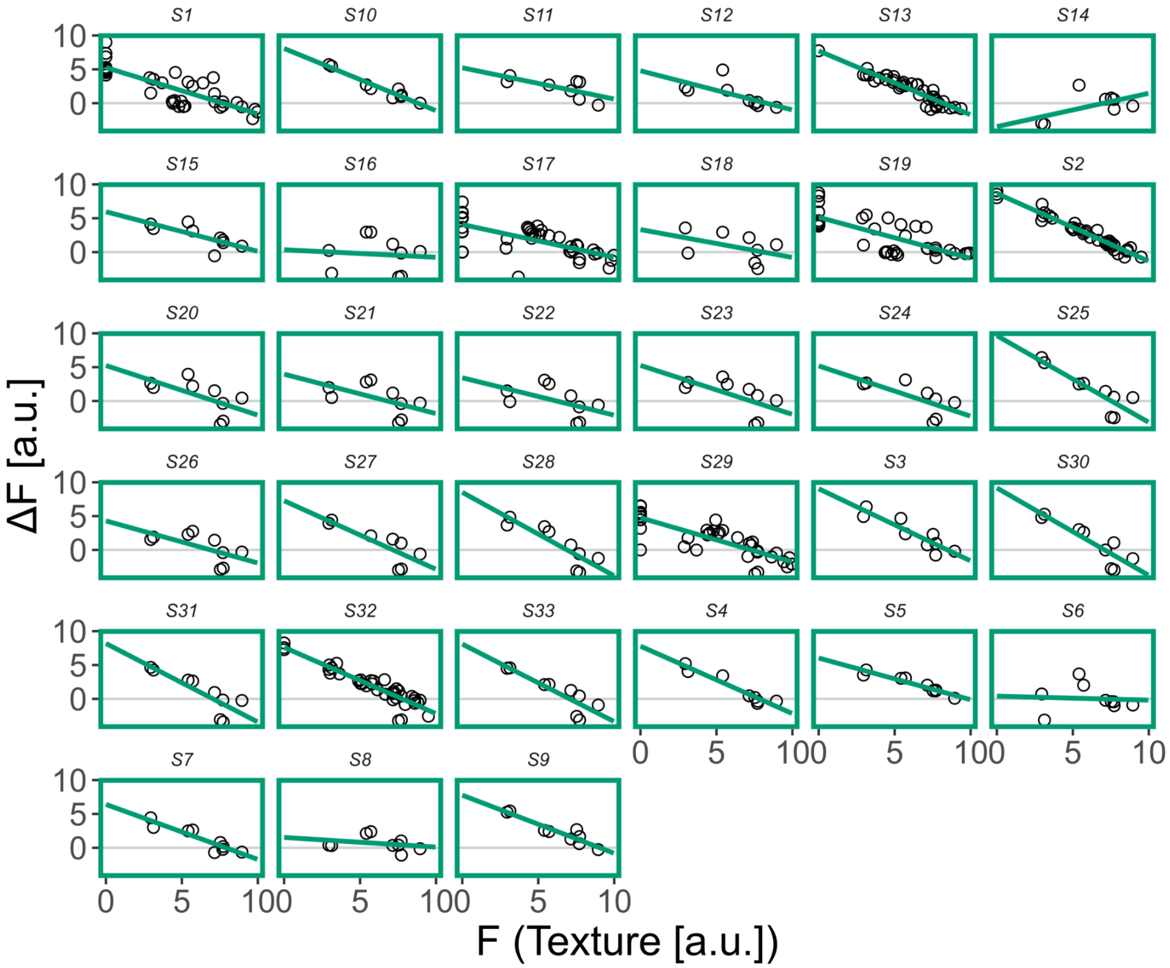


**Figure S2. Functional effect equations (FEEs) constructed for texture.** For each of the 33 assayed LAB strains, we have constructed FEEs based on the effect of each of the strains, with each strain assayed on at least 8 community backgrounds. The functional effect (ΔF) is defined as the difference in community function between the background alone and the background with the focal species.

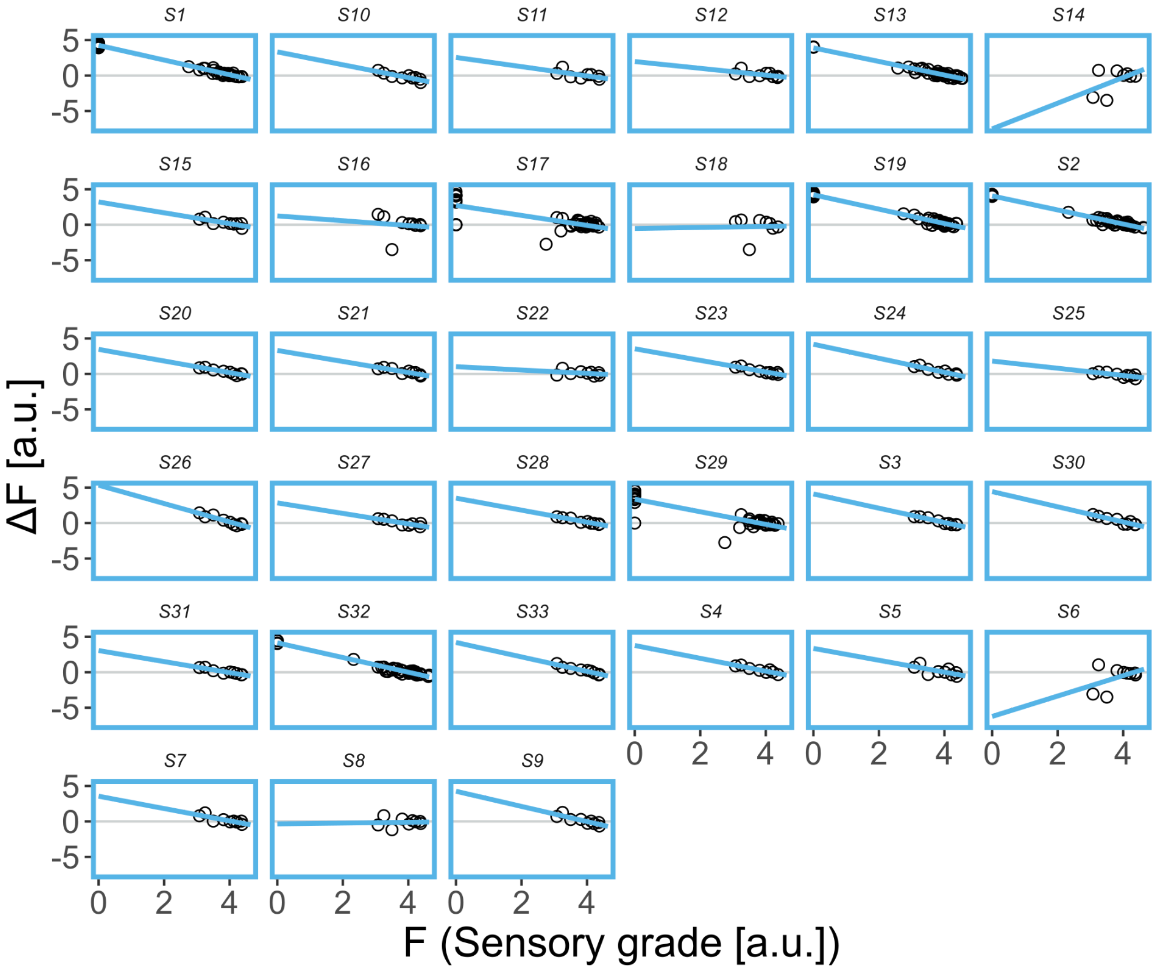


**Figure S3. Functional effect equations (FEEs) constructed for sensor grade.** For each of the 33 assayed LAB strains, we have constructed FEEs based on the effect of each of the strains, with each strain assayed on at least 8 community backgrounds. The functional effect (ΔF) is defined as the difference in community function between the background alone and the background with the focal species.

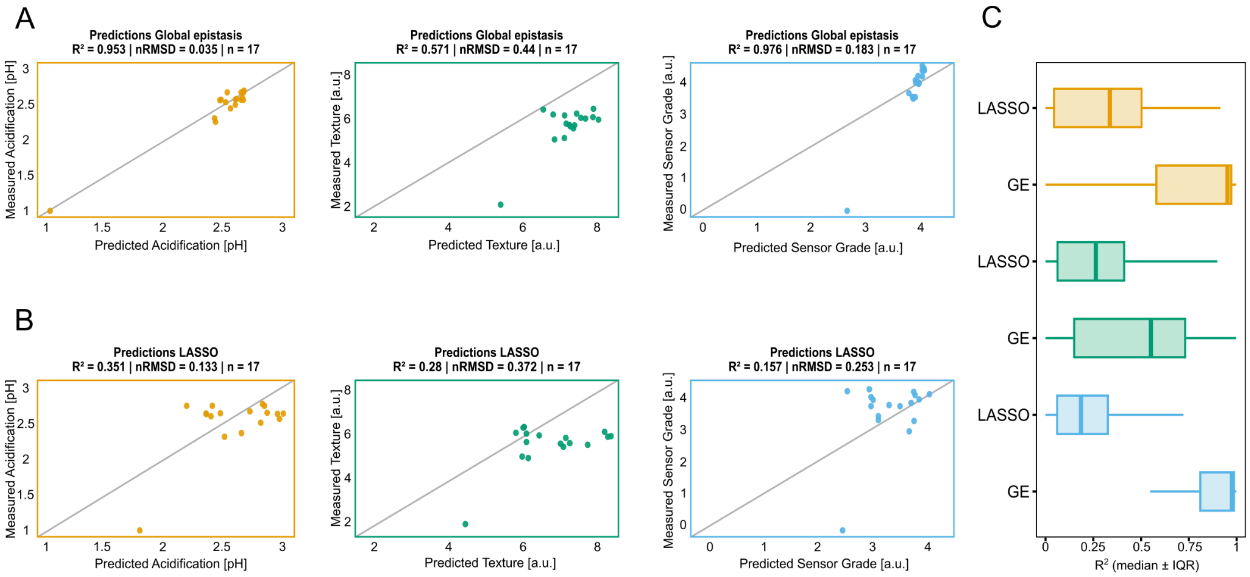


**Figure S4. Detailed comparison between FEE-based and LASSO predictions. A-B)** Predicted versus measured values using FEE stitching (A) or LASSO (B) on the validation set for the three functions (N = 17 in each case). Note that the panels in A reproduce the same coloured points shown in Fig. 2A (main text), represented here separately to allow direct comparison with LASSO predictions (panel B). **C)** Comparison of bootstrapped $R^2$ values (10000 iterations) between the FEE-stitching model and LASSO regression. Colors represent function, acidification (orange), texture (green) and sensor grade (blue). FEE-stitching consistently outperforms LASSO for the $R^2$ metric in all three functions (t-test $p<10^{-16}$ in all cases).

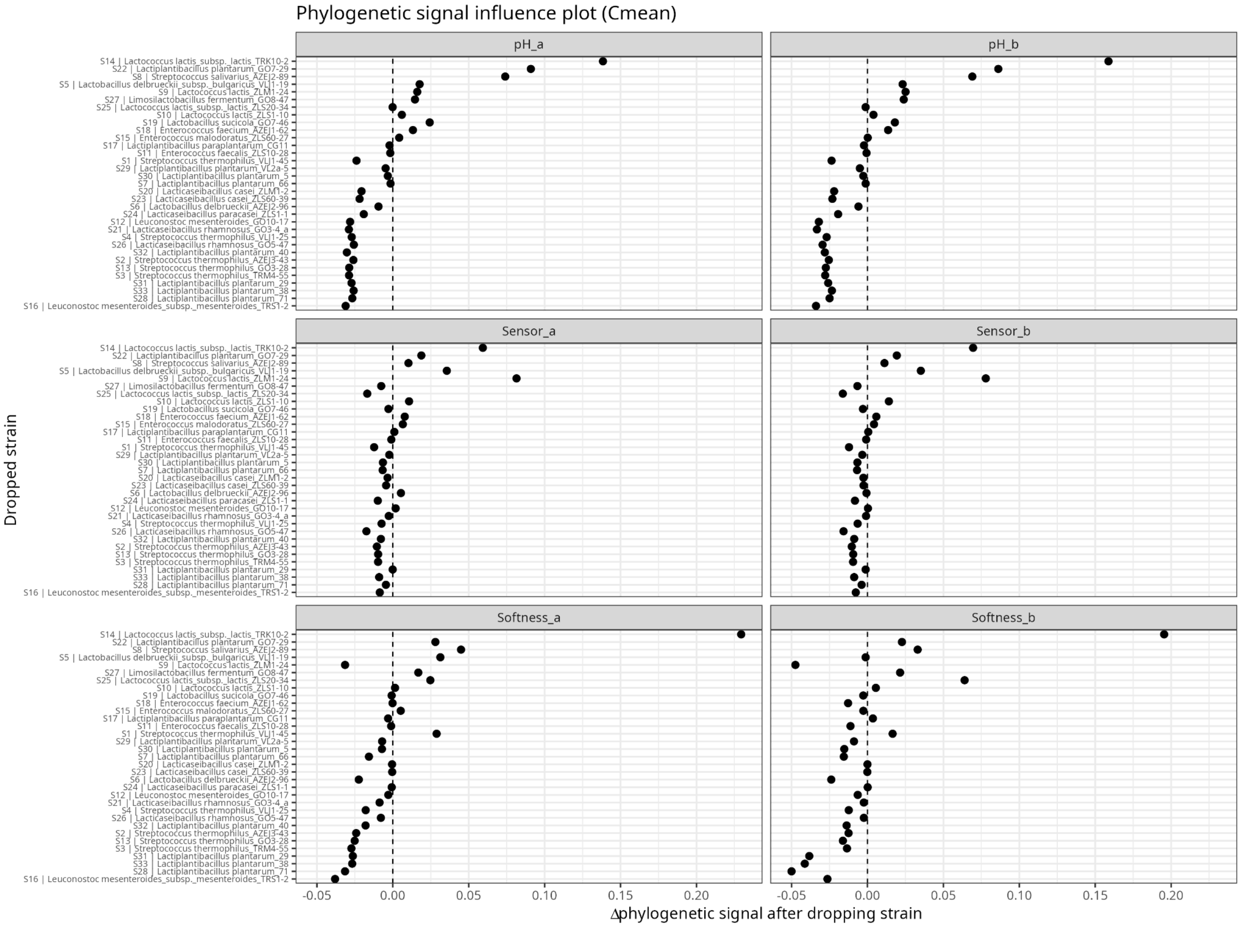


**Figure S5. Leave-one-out influence analysis of phylogenetic signal ($C_{mean}$) in FEE parameters.** Each panel corresponds to one FEE parameter (slope, *a*, or intercept, *b*) for acidification (pH), texture (Softness) and sensory grade (Sensory grade). Points show the change in phylogenetic signal, Δ = (signal estimated after removing the focal strain) − (signal estimated on the full 33-strain panel), with the signal measured as Abouheif's $C_{mean}$. Positive Δ indicates strains whose removal increases the signal (i.e. isolates departing from phylogenetic structure) whereas negative Δ indicates strains that reinforce phylogenetic trait structure. The dashed line marks Δ = 0. Strains are ordered along the y-axis by their mean Δ across all six parameters. Three isolates (S8, S14 and S22) show the largest positive Δ across multiple parameters and functions, identifying them as phylogenetically discordant; these were excluded from the conservation analysis ("pruned" tree, see main text).

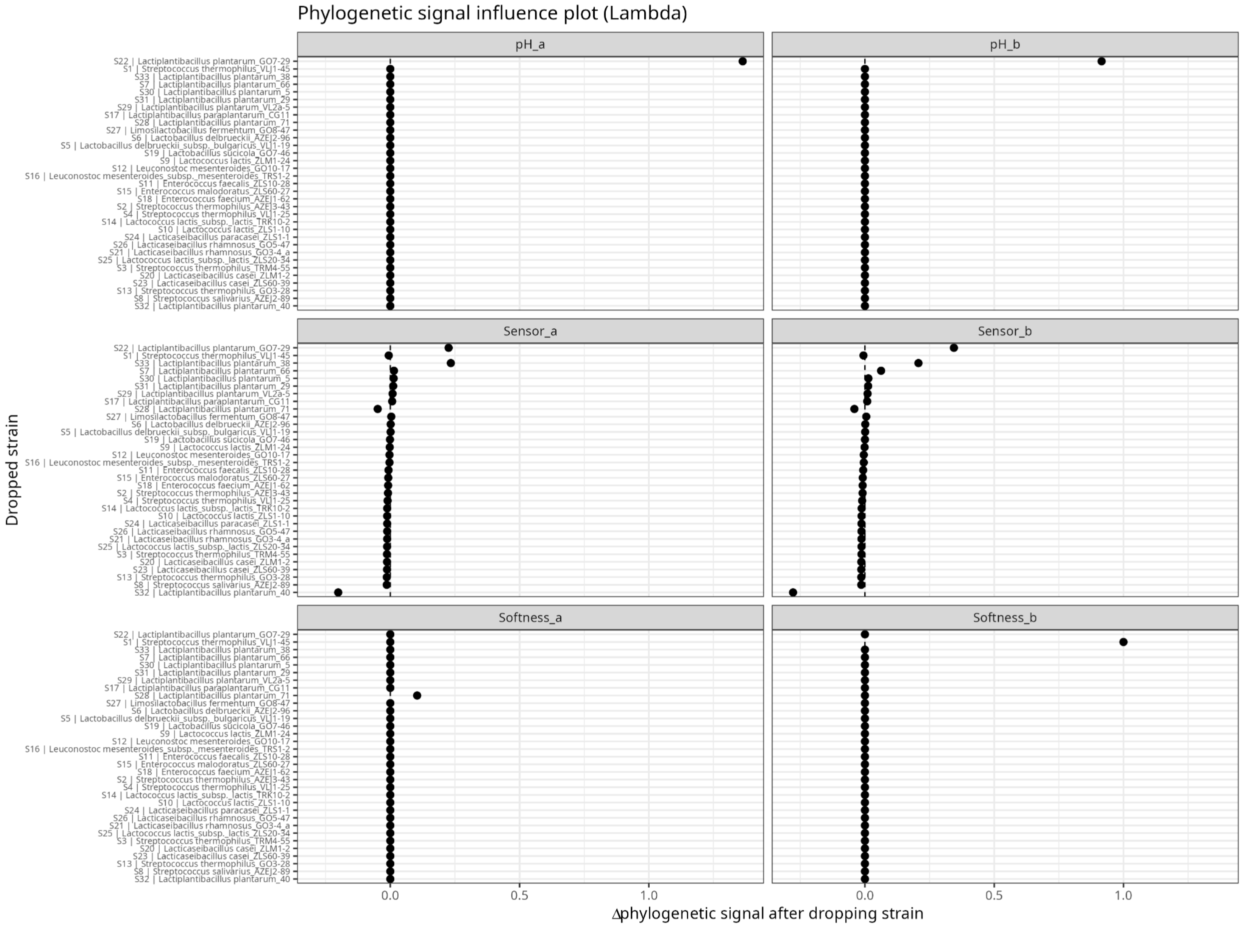


**Figure S6. Leave-one-out influence analysis of phylogenetic signal (Pagel's λ) in FEE parameters.** Figure analogous to Fig S5 for the Pagel's λ indicator.

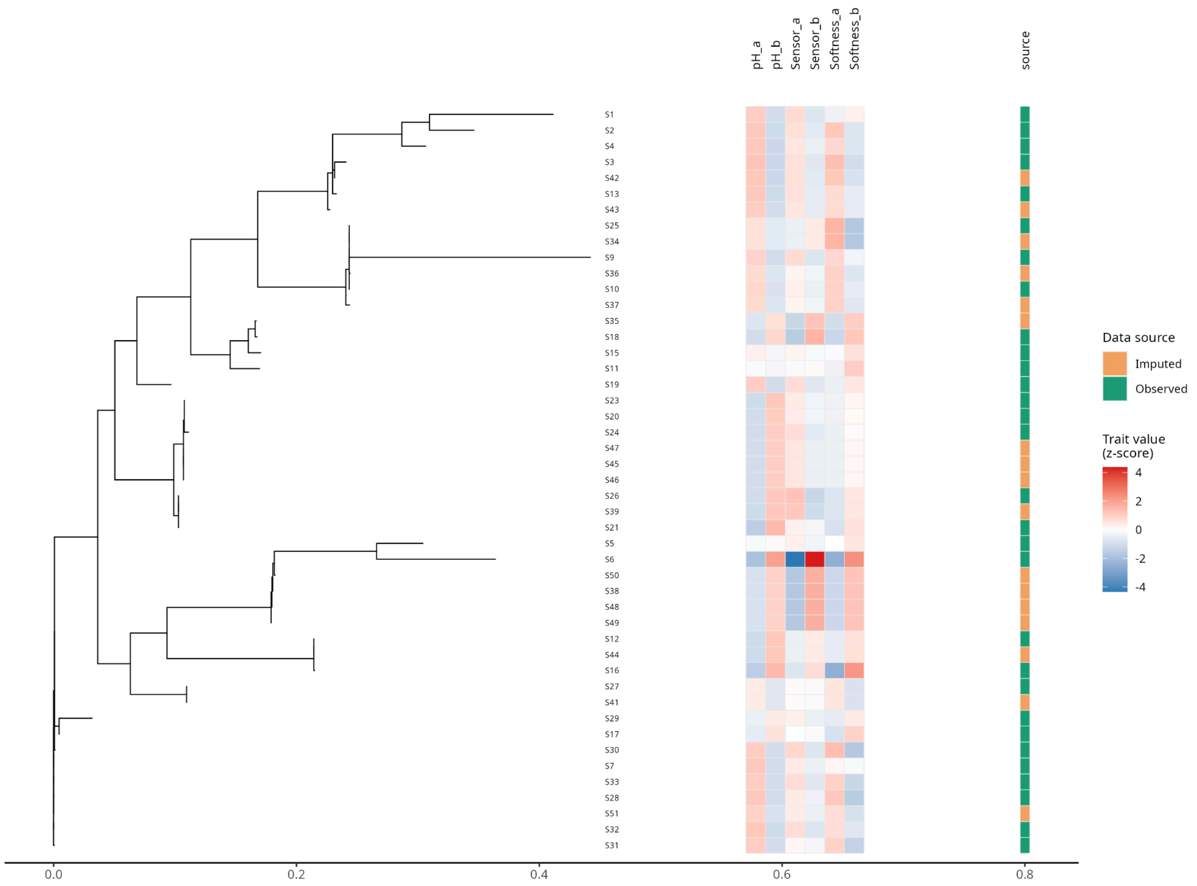


**Figure S7. Full phylogeny including imputed trait values for out of sample strains.** Phylogenetic tree analogous to Fig. 3A, with strains S8, S14 and S22 removed ("pruned" tree) and including the out-of-sample strains. The "source" column indicates whether a strain belongs to the initial pool ("Observed") or to the out-of-sample pool ("Imputed").

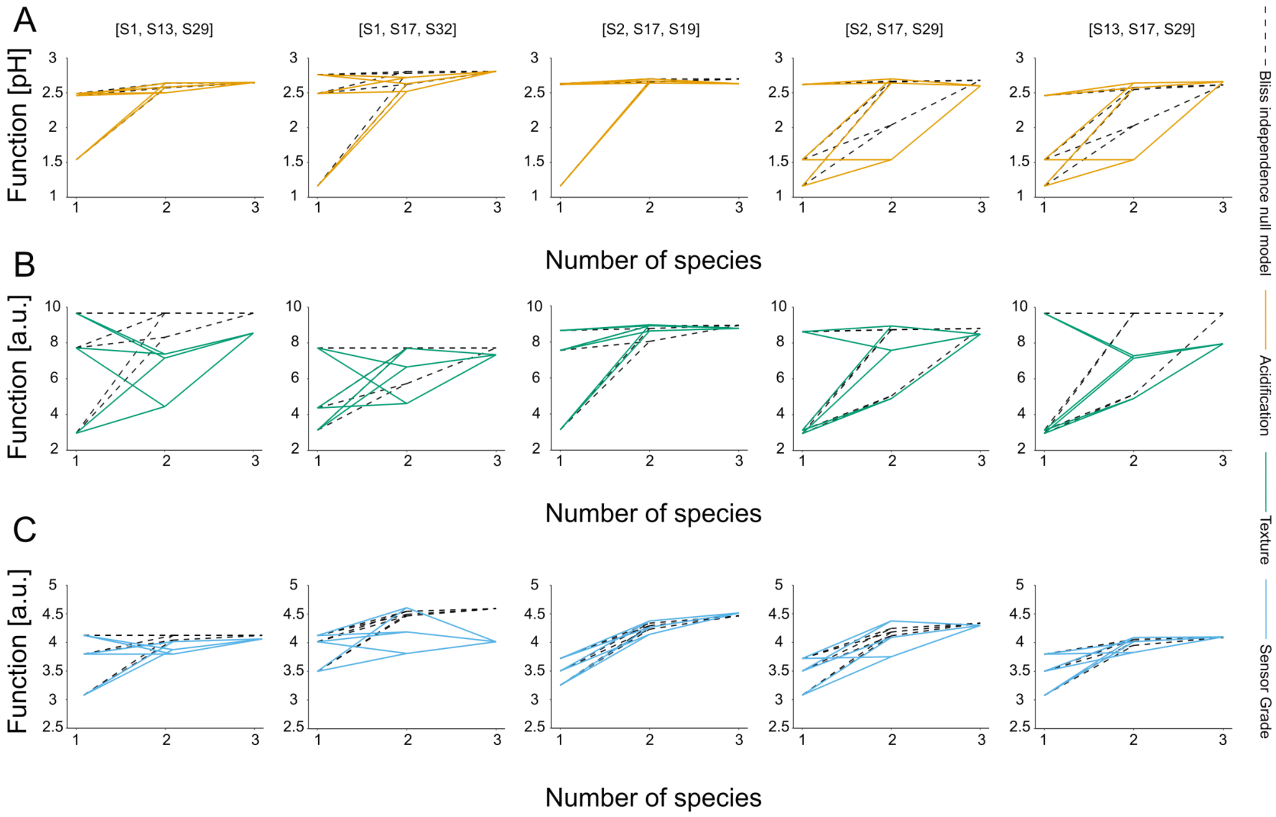


**Figure S8. Fully mapped combinatorial landscapes across measured functions.** (A-C) Functional landscapes for 5 out 8 subsets used for interaction coefficient calculations (with the remaining 3 shown in Fig. 4A, main text). Dashed black lines represent the predicted values of two- and three-member SynComs based on the Bliss independence null model (Methods), while coloured lines represent the measured functions for acidification (orange), texture (green) and sensor grade (blue). Deviations from the null model define interactions.

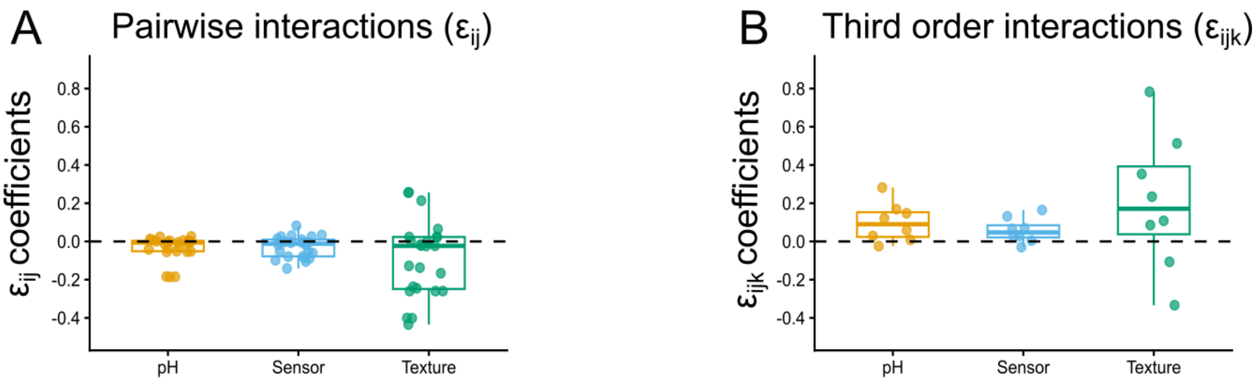


**Figure S9. A)** Non-absolute values of pairwise interaction coefficient, quantified as deviations from the Bliss null model across all community compositions for each of the eight subsets. **B)** Non-absolute values of third order interaction coefficients. Third-order interactions are defined as the deviation of the observed three-member community function from the value expected based on pairwise interactions among its constituent members.

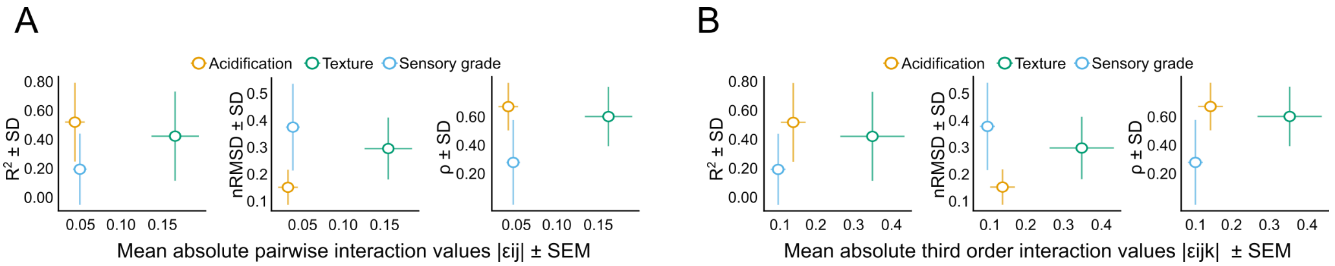


**Figure S10. A)** Relationship between mean absolute pairwise interaction strength and predictive performance for the out-of-sample set of SynComs (i.e. those including phylogenetically imputed FEEs). The x-axis shows the mean absolute pairwise interaction coefficient, while the y-axis shows predictive performance estimated by bootstrapping the out of sample set (N=14, 10000 iterations), quantified as $R^2$, nRMSD, and Spearman's ρ. Horizontal error bars indicate the standard error of the mean (SEM) of the absolute pairwise interaction coefficients, and vertical error bars indicate the standard deviation (SD) of the bootstrapped performance estimates. **B)** Relationship between mean absolute third order interaction strength and predictive performance for the out of sample set. The x-axis shows the mean absolute third order interaction coefficient, while the y-axis shows predictive performance estimated by bootstrapping the out of sample set (N=14, 10000 iterations), quantified as $R^2$, nRMSD, and Spearman's ρ. Horizontal error bars indicate the standard error of the mean (SEM) of the absolute third order interaction coefficients, and vertical error bars indicate the standard deviation (SD) of the bootstrapped performance estimates. In all plots, orange dots represent acidification, green texture and blue sensory grade.

| **Attribute Group** | **Description of Quality Levels / Defects** |
|---|---|
| **Appearance** | The soy yogurt should have a pale-yellow colour and a homogeneous appearance without visible whey separation. |
| (Visual impressions) | Excessively light or dark colour (deduct up to 1 point) |
| | Non-uniform colour (lighter and darker areas) (deduct up to 2 points) |
| | Excessive stringiness when scooped with a spoon (deduct up to 3 points) |
| | Visibly grainy appearance (deduct up to 2 points) |
| | Syneresis (whey separation) (deduct up to 3 points) |
| **Odor - Orthonasal** | The fermented soy drink should have a pleasant aroma of lactic acid and soy. |
| (Olfactory impressions) | Sharp odor (burning sensation in the nasal mucosa) (deduct up to 4.5 points) |
| | Aroma of an atypical acid (e.g., acetic, butyric) (deduct up to 3 points) |
| | Yeasty and/or foreign, atypical, unpleasant odor (e.g., animal fat, cardboard, etc.) (deduct up to 4.5 points) |
| | Excessively pronounced soy/pea aroma (deduct up to 3 points) |
| **Texture - Oral** | The product should have a homogeneous, cohesive mass. |
| (Palpatory impressions) | Excessively grainy/sandy texture (deduct up to 2 points) |
| | Non-homogeneous product (deduct up to 3 points) |

| | |
|---|---|
| | Product is either too liquid or too viscous (deduct up to 3 points) |
| | Excessively tough, rubbery, or sticky (deduct up to 2 points) |
| **Taste** | The fermented soy drink should have a pleasant, slightly sour taste, with a mild soy flavour. |
| (Gustatory impressions) | Excessively pronounced acidity (deduct up to 2 points) |
| | Presence of an atypical acid flavour (e.g., acetic, butyric) (deduct up to 3 points) |
| | Flavour of yeast, tallow, cardboard, scorched, etc. (deduct up to 4.5 points) |
| | Presence of bitterness and astringency (deduct up to 4 points) |
| | Excessively pronounced pea/soy flavour (deduct up to 3 points) |

**Supplementary table 1. Grading panel for Sensory grade function.** The table shows the criteria used to evaluate soy yoghurt quality. Fermented soy milk products were scored based on appearance, odor, oral texture and taste. For each category, desirable product characteristics and specific sensory defects are listed together with the corresponding point deductions applied during scoring.

| **Function** | **Community composition** | **nRMSD (mean ± SD)** | **$R^2$ (mean ± SD)** | **Spearman's ρ (mean ± SD)** | **N** |
|---|---|---|---|---|---|
| Acidification | **Imputed strains** | **0.11 ± 0.02** | **0.73 ± 0.26** | **0.69 ± 0.22** | **8** |
| | Imputed + assayed strain | 0.19 ± 0.12 | 0.72 ± 0.30 | 0.62 ± 0.35 | 6 |
| | Both | 0.16 ± 0.06 | 0.52 ± 0.27 | 0.65 ± 0.18 | 14 |
| Texture | **Imputed strains** | **0.29 ± 0.13** | **0.89 ± 0.16** | **0.81 ± 0.18** | **8** |
| | Imputed + assayed strain | 0.27 ± 0.09 | 0.24 ± 0.30 | 0.15 ± 0.56 | 6 |
| | Both | 0.30 ± 0.08 | 0.42 ± 0.30 | 0.58 ± 0.22 | 14 |
| Sensory grade | **Imputed strains** | **0.37 ± 0.18** | **0.50 ± 0.30** | **0.21 ± 0.48** | **8** |
| | Imputed + assayed strain | 0.31 ± 0.20 | 0.23 ± 0.27 | 0.34 ± 0.45 | 6 |
| | Both | 0.37 ± 0.13 | 0.20 ± 0.24 | 0.24 ± 0.31 | 14 |

**Supplementary Table 2. Bootstrapped prediction performance for the out of sample set.** The table summarizes prediction performance for the out of sample set containing only imputed strains, communities containing both imputed and assayed strains, and the combined set of both community types. Performance is reported for three measured functions: acidification, texture and sensory grade. For each function and community type, communities were reasampled with replacement for 10000 bootstrap iterations. Mean ± SD values are shown for nRMSD, $R^2$, and Spearman's rank correlation coefficient (ρ). The number of communities in each community type is indicated by N. Bolded rows correspond to values reported in Fig. 3C-E of the main text.

**Supplementary Table 3 (provided as supplementary file in excel: "Supplementary table 3 - strain annotation + 16s sequences"). Strain notation and taxonomy.** The table lists the strain notation IDs used throughout the study, the corresponding taxonomic names, and the associated 16S rRNA gene sequences. Strains denoted S1-S33 refer to the assayed strains, whereas S34-S51 refer to strains with phylogenetically imputed FEEs.

**Supplementary Table 4 (provided as supplementary file in excel: "Supplementary table 4 - Community annotation"). Strain composition of tested communities.** The table lists the community notation IDs used throughout the study, the corresponding strain composition, and the community type. Community notation IDs starting with C refer to communities used in the training dataset (N = 307), IDs beginning with CV refer to validation set communities (N=17) and IDs beginning with COS refer to out-of-sample communities (i.e. those including strains with phylogenetically imputed FEEs, N=14).
